# Artifact-guided restoration improves visualization of cellular ultrastructure in cryo-FIB-SEM volumes

**DOI:** 10.64898/2026.11.09.765396

**Authors:** Ningyan Zhang, Lydia-Marie Joubert, Gong-Her Wu, Guan-Chin Su, Pingting Liu, Manjari Mishra, Pengyang Li, Jorge Sanz-Ros, Shirit Einav, Bo Wang, Inma Cobos, Min Xu, Muyuan Chen

## Abstract

Cryogenic focused ion beam scanning electron microscopy (cryo-FIB-SEM) enables volumetric imaging of vitrified specimens, but charging artifacts, high noise and weak membrane contrast hinder the interpretation of biological features. Here we present a two-stage deep-learning-based framework that translates cryo-FIB-SEM images toward the domain of conventional room-temperature FIB-SEM (RT-FIB-SEM), suppressing large-scale charging artifacts and enhancing local membrane signal. Held-out synthetic tests show improved agreement with constructed reference targets, while results on four experimental cryo-FIB-SEM datasets show reduced visible artifacts and clearer membrane features. Processed cryo-FIB-SEM volumes from our two-stage model can be directly analyzed by tools designed for RT-FIB-SEM, and an existing pretrained segmentation model is applied to our model output, yielding extensive predictions of membrane structures. This post-acquisition approach improves the visual inter-pretability and downstream analysis capabilities of cryo-FIB-SEM data without modifying imaging hardware.

## 1 Introduction

Focused ion beam–scanning electron microscopy (FIB-SEM) enables 3D imaging of cellular ultrastructure through successive ion-beam milling and scanning electron microscopy image acquisition. Advances in RT-FIB-SEM have made it possible to acquire large volumes with nearly isotropic nanometre-scale sampling, revealing the organization of cells, organelles and tissues [1, 2]. Complementary advances in automated segmentation have enabled systematic annotation and quantitative analysis of organelles within these datasets [3]. These high-quality volumes provide both biological insight and a source of structural information for developing learning-based image analysis methods.

However, conventional RT-FIB-SEM preparation of biological specimens typically involves chemical fixation, dehydration, heavy-metal staining and resin embedding, procedures that can alter the hydrated cellular environment. By contrast, cryo-FIB-SEM images vitrified specimens, preserving cellular architecture in a near-native hydrated state and enabling complementary 3D and correlative studies [4–6]. This advantage comes with imaging challenges: low intrinsic contrast and charge accumulation in poorly conductive specimens can obscure ultrastructure [5, 7]. Charging-associated dark regions and directional streaks, together with uneven intensity backgrounds, noise and loss of focus, can further compromise the annotation of membranes [5, 8, 9].

Developments in cryo-FIB-SEM data collection methods, including low-voltage imaging and high-efficiency detection, have alleviated some of these limitations. Cryo-plasma FIB-SEM workflows have extended 3D imaging to diverse vitrified biological specimens [5], while interleaved scanning can attenuate charging and reveal features that would otherwise remain masked [7]. Post-acquisition processing complements these acquisition-side advances by targeting residual artifacts in previously acquired volumes without requiring changes to imaging hardware or acquisition protocols.

Existing computational methods address image registration, charging mitigation, artifact correction, denoising and enhancement in cryo-FIB-SEM and related volume electron microscopy (vEM) data [8–11]. However, charging-related distortions, stripes, uneven backgrounds, noise and loss of focus can coexist across spatial scales and vary between adjacent slices, complicating joint correction and membrane preservation [5, 6, 8]. Paired degraded and artifact-free cryo-FIB-SEM images are also rarely available to supervise restoration.

Here we present a two-stage post-acquisition framework that translates cryo-FIB-SEM images toward the domain of RT-FIB-SEM, with reduced imaging artifacts and enhanced membrane contrast. We construct stage-specific supervised training pairs from these structural references, incorporating experimental cryo-FIB-SEM charging patterns alongside complementary synthetic degradations. Results of the approach are evaluated on held-out synthetic tests and four acquired cryo-FIB-SEM datasets, demonstrating its enhancement of image quality on a range of specimens. Additionally, we show the method’s capability in the cryo-to-RT domain translation by applying an existing pretrained membrane segmentation model [12], designed for RT-FIB-SEM data, to cryo-FIB-SEM volumes processed by our models.

## 2 Results

### 2.1 Task-specific design of the two-stage framework

The framework comprises two independently trained models for artifact suppression and membrane contrast enhancement (Fig. 1). Both models use RT-FIB-SEM data and stage-specific synthetic degradations for supervision, avoiding the need for paired degraded and artifact-free cryo-FIB-SEM images. We further examined whether the processed cryo-FIB-SEM volumes could support downstream analysis using an existing membrane-segmentation model developed for RT-FIB-SEM data.

**Fig. 1.**
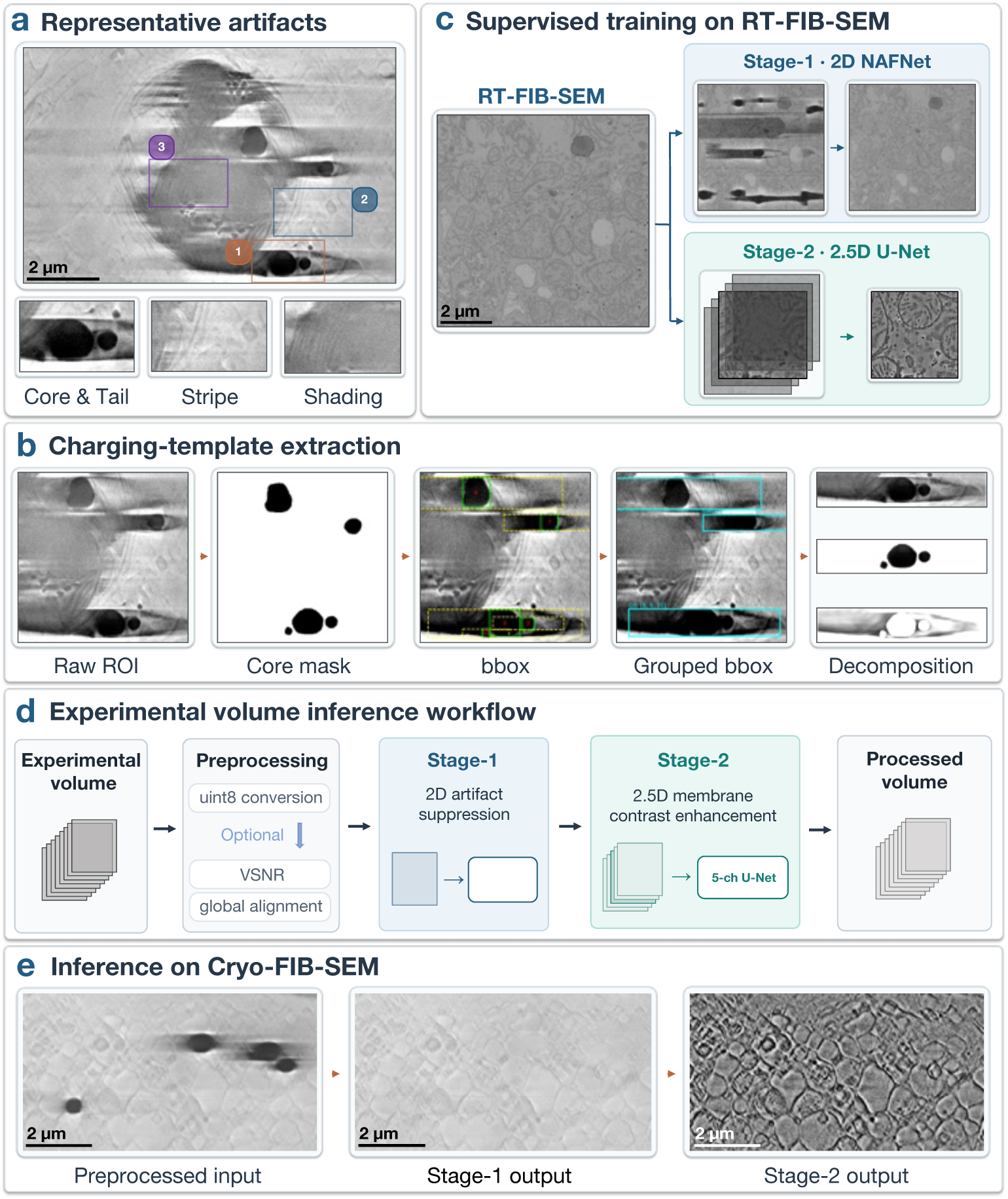
Two-stage artifact suppression and membrane contrast enhancement in cryo-FIB-SEM. a, Representative charging, stripe/bundle and shading artifacts in experimental cryo-FIB-SEM images. **b,** Extraction and decomposition of charging-artifact templates from cryo-FIB-SEM data. **c,** Separately constructed supervised training pairs for Stage-1 (2D) and Stage-2 (2.5D), both derived from RT-FIB-SEM data. **d,** Workflow on experimental data comprising input preparation, Stage-1 artifact suppression and Stage-2 membrane contrast enhancement. **e,** Representative views of primary human small airway epithelial cells (HSAEC), showing the preprocessed input, Stage-1 output and Stage-2 output. Scale bars, 2 *µ*m (a, c and e).

The two stages use different spatial contexts to address their respective tasks. Stage-1 targets structured high-contrast, large-scale artifacts (Fig. 1a) [5, 8]. Although individual artifacts may occupy only part of a slice, their lateral extent can be substantial. We therefore trained a 2D NAFNet-based model [13] on 1,024×1,024-pixel images, providing a large field of view for artifacts that could be truncated by smaller training crops. Membrane structures can exhibit meaningful correspondence across adjacent slices, but imaging artifacts may vary substantially between them. Processing each slice independently allows Stage-1 to exploit the artifact information present within that image without relying on potentially inconsistent inter-slice artifact patterns or requiring their joint synthesis across a stack.

Stage-2 instead focuses on membrane contrast enhancement and denoising throughout the image, rather than on the suppression of large-scale patterns. In this case, neighboring slices can provide complementary structural context, especially when the biological features in the central slice are corrupted by noise, local artifacts or loss of focus. Accordingly, Stage-2 uses a five-channel U-Net-based model [14] that takes 256×256-pixel patches from five consecutive slices and predicts a membrane-enhanced central patch (Fig. 1c). The smaller spatial field allows training across local regions and supports center-referenced patch alignment during experimental data inference. The model is trained to use neighboring information despite slice-dependent corruption and local displacement, rather than assuming exact correspondence between slices. Suppression of residual scan-direction stripes remaining after Stage-1 is included as a secondary objective.

Both stages use the publicly available RT-FIB-SEM mouse-liver volume as their source of biological features [2], but their synthetic inputs and supervision targets are constructed independently (Fig. 1c). For Stage-1, source images are adapted to four background-intensity levels informed by acquired cryo-FIB-SEM data. They are then combined with charging patterns derived from manually annotated cores and associated directional intensity residuals in acquired cryo-FIB-SEM images (Fig. 1b), together with additional parameterized intensity artifacts. Maintaining the contrast relationship between each artifact and its surrounding background is important for supervising intensity-sensitive correction. Stage-2, by contrast, uses separately synthesized degradations of five-slice patch inputs paired with membrane-enhanced central-patch targets. Before patch extraction, the Stage-2 training slices undergo full-slice intensity preprocessing to suppress low-frequency backgrounds and emphasize local structural information. These distinct input representations reflect the different objectives of artifact suppression and membrane contrast enhancement, rather than differences in the biological source data.

For inference on cryo-FIB-SEM data, volumes are first converted to the required input format. Preprocessing steps, including variational stationary noise removal (VSNR) for curtaining suppression [15], global translation-only alignment, and cropping, can be selectively applied depending on the characteristics of each dataset (Fig. 1d). Stage-1 then suppresses prominent artifacts independently in each slice. Stage-1 outputs undergo full-slice intensity preprocessing before Stage-2 patch extraction. Before Stage-2 prediction, the four neighboring patches are aligned locally to the central patch to reduce residual lateral displacement. Stage-2 then predicts overlapping central patches from five-slice inputs, which are combined into an output slice without changing the acquired axial sampling. Taken together, the two-stage models remove the artifacts and enhance membrane contrast when applied to acquired cryo-FIB-SEM datasets (Fig. 1e).

### 2.2 Learning to suppress large-scale imaging artifacts

We evaluated Stage-1 artifact suppression using held-out synthetic image pairs derived from RT-FIB-SEM structural references (Fig. 2a). These controlled comparisons allowed us to assess agreement with constructed supervision targets and preservation of artifact-free image content.

**Fig. 2.**
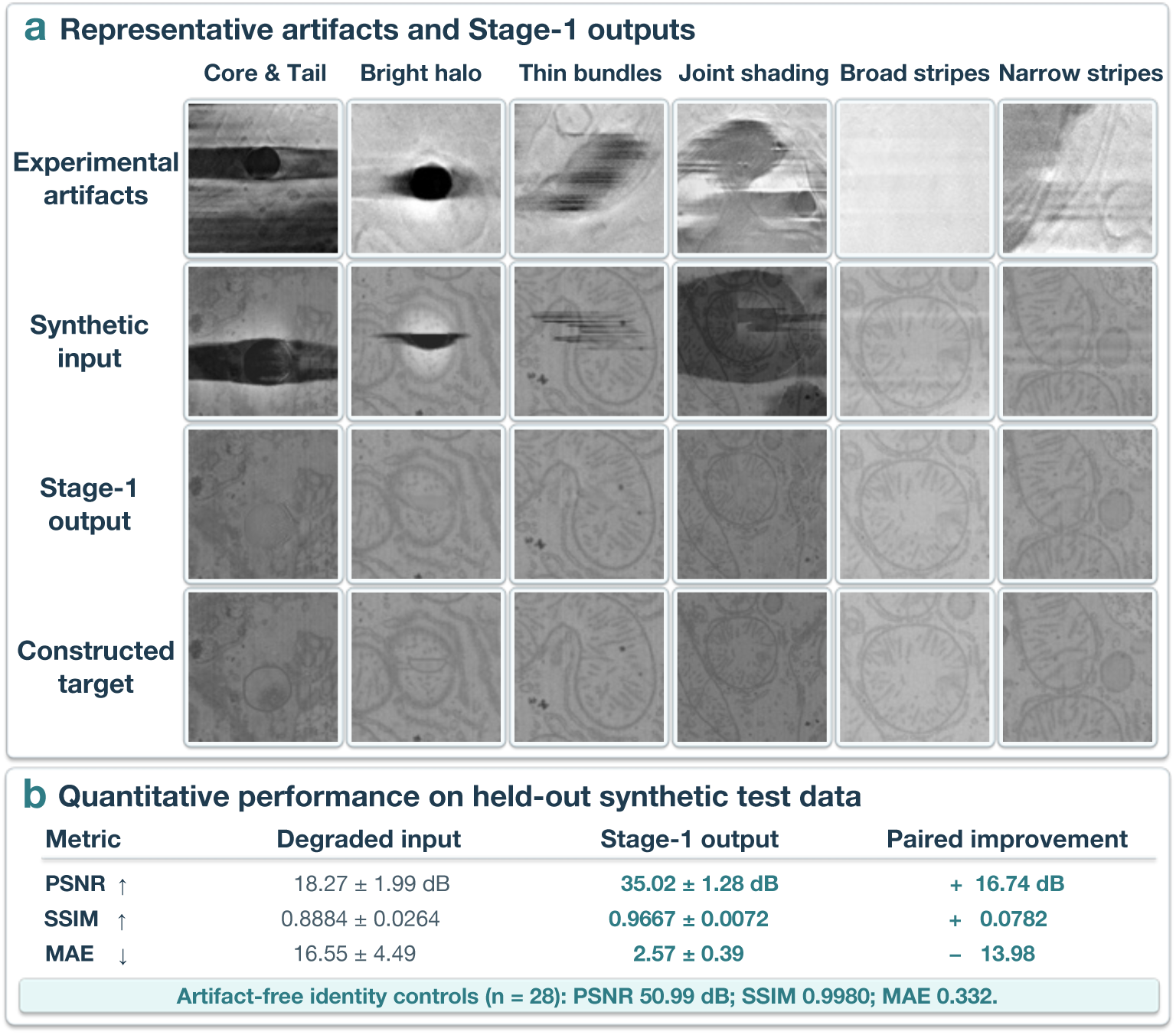
Synthesis guided by experimental artifacts and Stage-1 artifact suppression. a, Six representative artifact types illustrated by experimental cryo-FIB-SEM examples (top row), synthetic degraded inputs, Stage-1 outputs and constructed reference targets (bottom three rows). **b,** Peak signal-to-noise ratio (PSNR), structural similarity index (SSIM) and mean absolute error (MAE) on held-out synthetic test images (*n* = 164), comparing degraded inputs and Stage-1 outputs with their reference targets. Values are mean *±* standard deviation; paired improvements are mean within-image changes. Artifact-free identity tests (*n* = 28) are reported separately.

Stage-1 training incorporated six representative artifact appearances: charging cores with dark directional tails, bright halos, thin line bundles, joint shading, broad bright stripes or seams, and alternating narrow stripes (Fig. 2a). Stripe and bundle patterns were modeled parametrically, whereas irregular charging-related distortions were synthesized using empirical templates. We extracted charging-core masks and directional intensity residuals from a region of one cryo-FIB-SEM dataset spatially separated from the regions used for experimental-data evaluation. These data informed artifact synthesis and background-intensity adaptation but did not contribute biological structural supervision, which was provided exclusively by the RT-FIB-SEM mouse-liver references [2]. Charging-derived fields also informed halo and broad-shading synthesis. Multiple artifact types could be introduced into the same synthetic image to reflect the diversity of degradation patterns encountered in practice.

In the clean supervision targets, pixels inside the synthetic charging-core masks were replaced with background-matched texture and a narrow peripheral intensity cue, while the original structural references were retained outside the masks. This construction avoids treating obscured membrane structures beneath charging cores as known ground truth. Training additionally emphasized strong intensity deviations, transitions between corrections of opposite polarity, and weak but extended shading, while artifact-free identity examples discouraged unnecessary changes to unaffected image content.

On 164 held-out synthetic test images (1,024 × 1,024 pixels), Stage-1 substantially improved agreement with the constructed references, with a mean paired PSNR gain of 16.74 dB and corresponding improvements in SSIM and MAE (Fig. 2b). Representative Stage-1 outputs show suppression of charging-related artifacts, stripes and shading, while the 28 artifact-free identity tests showed minimal changes to the input images (Fig. 2a,b). These quantitative results characterize performance against constructed synthetic targets and do not directly establish artifact-suppression accuracy on acquired cryo-FIB-SEM data.

### 2.3 Learning to enhance membrane contrast using multi-slice context

We evaluated Stage-2 membrane contrast enhancement and denoising using controlled synthetic degradations of held-out RT-FIB-SEM mouse-liver data, using computationally enhanced central-slice images as reference targets (Fig. 3a).

**Fig. 3.**
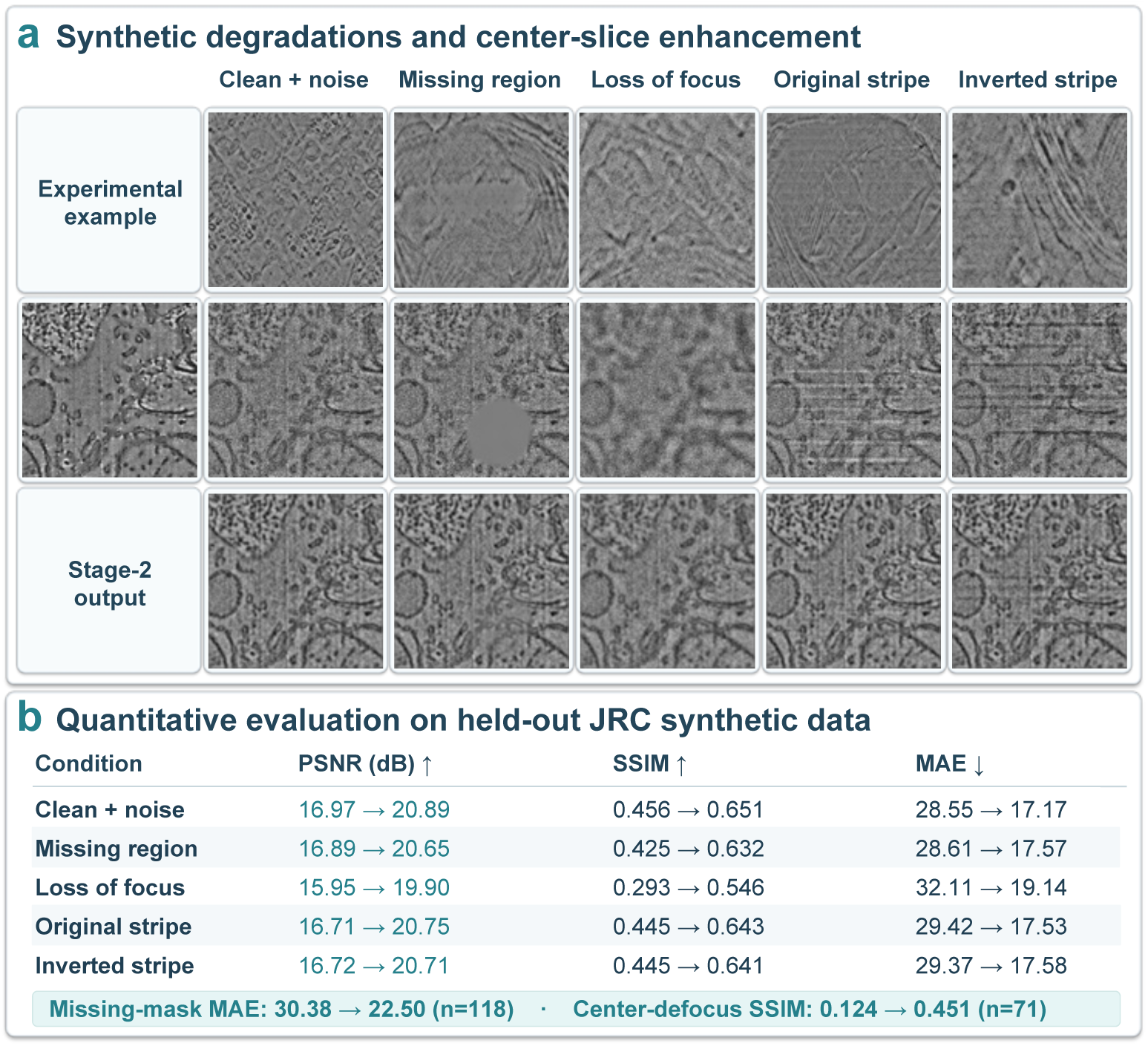
Synthetic degradations and Stage-2 membrane contrast enhancement. a, Unpaired cryo-FIB-SEM examples (top), synthetic degradations (middle) and Stage-2 outputs (bottom) across five conditions. A shared RT-FIB-SEM mouse-liver source region is shown at middle left. The central-slice views are display crops, not the 256 *×* 256 network input patches. **b,** PSNR, SSIM and MAE for degraded central patches and Stage-2 outputs against computationally enhanced reference targets on held-out data. Visualized examples emphasize central-slice degradation, whereas quantitative evaluation includes the full set of 144 patches per condition.

All Stage-2 synthetic training inputs contained Gaussian noise in the processed-intensity domain. Further perturbations included missing regions applied to selected patches within each five-slice patch input, defocus affecting a single source slice, and spatial shifts in neighboring patches. Residual scan-direction stripes of either intensity polarity were also included to address interference remaining after Stage-1. The U-Net combined the five-slice context while retaining a direct full-resolution pathway from the central input patch as the spatial reference for prediction.

Across the five displayed conditions, Stage-2 reduced noise and enhanced the visibility of membrane-associated boundaries relative to the degraded central inputs (Fig. 3a). In synthetic examples where the central input patch contained missing signal or was affected by loss of focus, Stage-2 outputs showed improved local structural contrast. Residual stripes of both polarities were also attenuated as a secondary artifact-suppression objective. Quantitative evaluation on 144 matched patches per condition from three held-out blocks showed improvements across all five displayed degradation conditions (Fig. 3b). Mean PSNR gains ranged from 3.76 to 4.03 dB, accompanied by increased SSIM and reduced MAE. For the 118 patches with missing regions affecting the central input patch, missing-mask MAE decreased from 30.38 to 22.50; for the 71 central-slice defocus cases, SSIM increased from 0.124 to 0.451.

### 2.4 Artifact suppression and membrane contrast enhancement in acquired cryo-FIB-SEM data

We next evaluated the trained two-stage models on four acquired cryo-FIB-SEM datasets representing distinct biological samples and a range of artifact conditions (Fig. 4). The publicly available *Chlamydomonas reinhardtii* dataset (EMPIAR-11275) contains well-resolved ultrastructure with comparatively limited charging [16], whereas the three in-house datasets exhibit more pronounced and varied intensity distortions. These evaluations examined whether the two-stage framework could suppress artifacts and enhance membrane contrast across experimental datasets without dataset-specific model retraining. All four datasets were processed using the same two-stage inference procedure, including full-slice intensity preprocessing and center-referenced local alignment before Stage-2 prediction. Optional intensity-and Fourier-statistics matching was enabled for the results shown here (Methods 4.8). The fields of view in the figures are crops from the processed full-slice results, not individual Stage-2 input patches; restored slices are assembled from overlapping predictions based on five-slice 256 × 256 patch inputs (Methods 4.8).

**Fig. 4.**
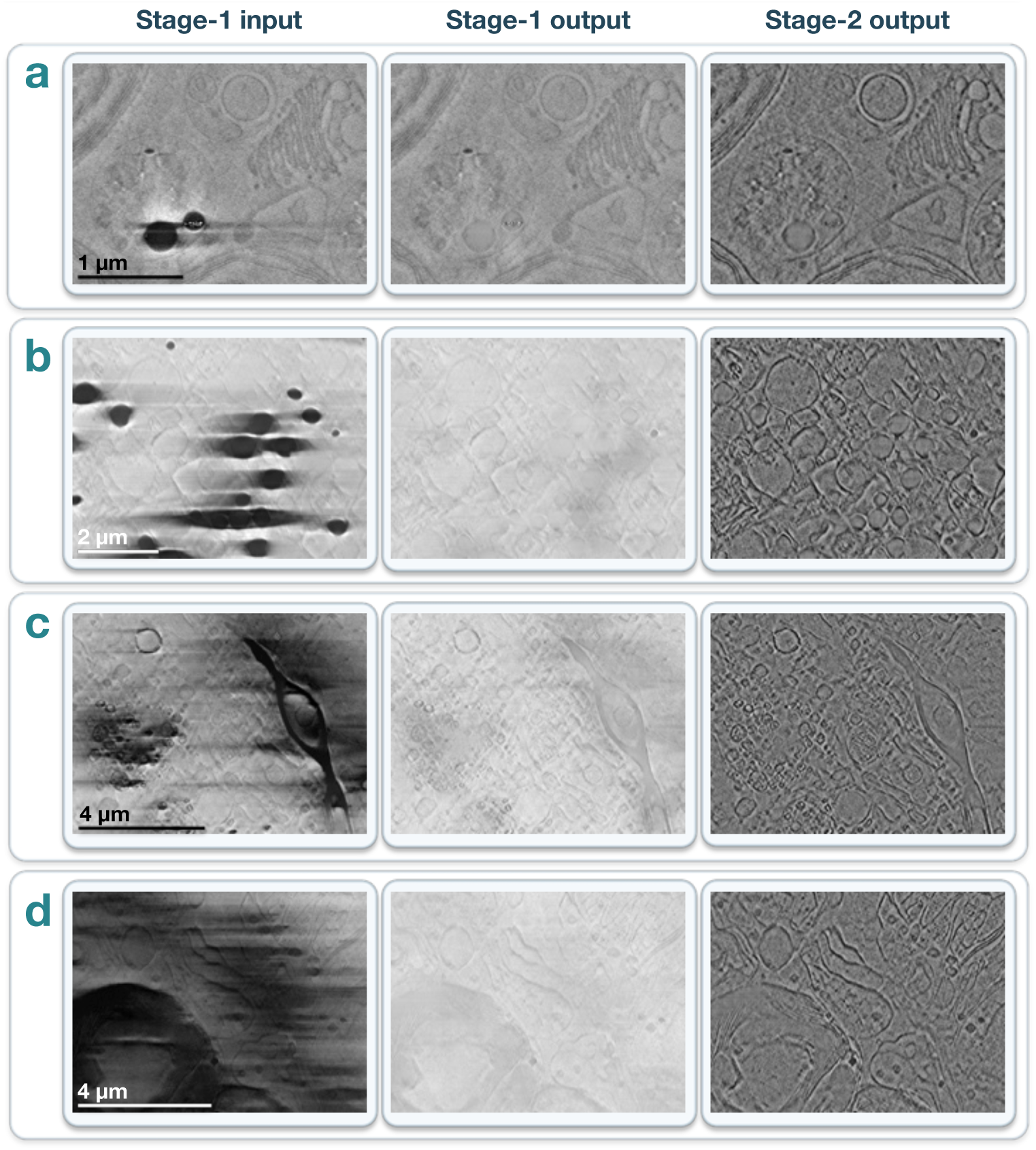
Two-stage processing of experimental cryo-FIB-SEM datasets. a, *Chlamydomonas reinhardtii*. **b,** Primary human small airway epithelial cells (HSAEC). **c,** Mouse brain tissue. **d,** Worm–algae symbiotic system. For each sample, the three columns show the Stage-1 input, Stage-1 output and Stage-2 output (left to right) for the same field of view. Scale bars are shown in the Stage-1 input images and apply to all three images in each row: 1 *µ*m (a), 2 *µ*m (b) and 4 *µ*m (c,d).

The relatively clean *Chlamydomonas* dataset [16] provides a useful contrast to more severely affected acquisitions (Fig. 4a). Stage-1 attenuates limited localized charging artifacts, while Stage-2 increases the contrast of fine membrane features. We further examined a structure-free ice region and a structure-poor biological region to qualitatively assess the generation of conspicuous membrane-like features in areas with limited visible structural evidence (Supplementary Fig. 1).

The second dataset, the primary human small airway epithelial cells (HSAEC), presents a substantially more challenging intensity landscape (Fig. 4b), with numerous dark charging cores, directional tails and adjacent bright distortions extending across the field. Stage-1 reduces the prominence of these large-scale artifacts, and Stage-2 enhances the contrast of the lipid membranes, facilitating identification of target organelles within the tissue.

The next example shows brain tissue from an App^SAA^ knock-in mouse [17] (Fig. 4c), containing localized lipid-rich regions associated with strong charging artifacts, including scan-direction stripes and irregular shading. The successive model outputs distinguish the reduction of long-range intensity disruptions from the subsequent enhancement of local membrane contours. We also examined structural consistency across five consecutive slices of the mouse brain volume (Supplementary Fig. 5).

The final and most challenging example comprises a symbiotic system in which a marine acoel worm, *Convolutriloba longifissura*, hosts green algae *Tetraselmis* [18]. The algal cells reside within animal tissue in between animal cells and can accumulate substantial lipid reserves that may contribute to host metabolism. The displayed region exhibits strong, often overlapping artifacts, including charging cores with tails, stripes and spatially varying background intensity (Fig. 4d). Stage-1 suppresses much of the broad interference, although the intermediate image retains weak membrane contrast; Stage-2 then reveals more clearly defined membrane features across the image, improving visual distinction between algal and animal cells. We additionally compared our framework with the artifact-correction methods implemented in Okapi-EM and SEM3De on the same region (Supplementary Fig. 4).

Together, these examples illustrate complementary changes produced by the two stages across distinct experimental conditions, rather than restoration of a single isolated artifact morphology. Larger-field views of the same slices presented in Fig. 4 are provided for all four datasets (Supplementary Figs. 6–9).

### 2.5 Membrane segmentation of processed cryo-FIB-SEM volumes

vEM datasets contain rich biological information, and many computational methods have been developed for conventional vEM that support tasks including membrane segmentation, organelle detection, and reconstruction of cellular connectivity [3, 12, 19]. Rather than developing separate analysis tools for cryo-FIB-SEM data, we investigated whether cryo-FIB-SEM volumes processed by our two-stage framework could be directly analyzed using existing methods designed for conventional vEM.

Here we used the task of membrane annotation as an example and applied the pretrained CebraNET membrane predictor [12, 20] to corresponding original and Stage-2-processed volumes without retraining or fine-tuning. Within each dataset, model weights, axial resampling settings and the membrane-probability threshold (0.30) were identical for original and restored inputs; predictions were mapped back to the acquired slice coordinates.

First, we compared membrane predictions by overlaying the CebraNET binary masks on corresponding 2D input images (Fig. 5a,b). In the worm-algae symbiosis dataset, extensive overlapping artifacts obscure weak membrane signals in the original data, producing broad artifact-associated predictions and incomplete membrane contours (Fig. 5a). With Stage-2-processed inputs, membrane predictions more closely follow visible cellular boundaries, including the densely stacked thylakoid membranes of algal chloroplasts. The corresponding 3D rendering illustrates the spatial organization of predicted membrane surfaces. In the mouse brain example, strong intensity disturbances coincide with broad artifact-associated predictions and incomplete membrane boundaries in the original input (Fig. 5b). The Stage-2-processed volume yields more detailed predictions along visible cellular and organelle membranes. The 3D visualization shows predicted membrane surfaces surrounding vesicle-like structures, together with manually highlighted mitochondria.

**Fig. 5.**
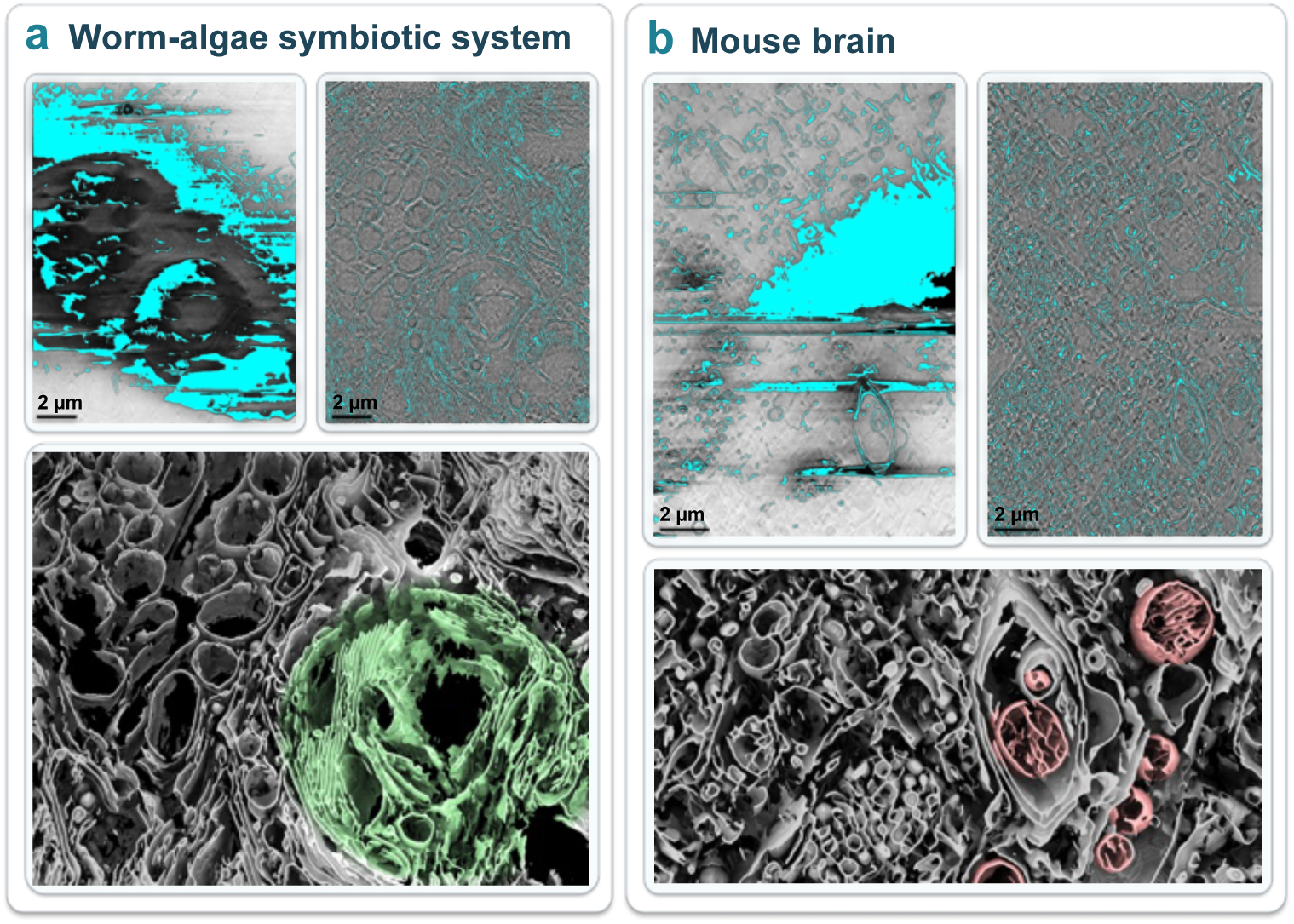
Membrane prediction and 3D visualization in processed cryo-FIB-SEM volumes. a, Worm-algae symbiotic system. **b,** Mouse brain. In each panel, the upper images show CebraNET membrane predictions (cyan) overlaid on corresponding preprocessed Stage-1 inputs (left) and Stage-2-processed slices (right). The lower images show 3D surface renderings of predicted membranes from the Stage-2-processed volumes. The algal region (green, a) and mitochondria (pink, b) were manually annotated to highlight representative biological structures. Scale bars, 2 *µ*m (upper 2D overlays only).

Taken together, these qualitative comparisons indicate that our two-stage model can make existing data analysis tools designed for conventional vEM more applicable for the noisy and artifact-corrupted cryo-FIB-SEM datasets. While the differences between the original and restored results are visually clear, quantifying the improvement in membrane segmentation remains challenging without established ground truth. However, manual annotation of conventional vEM datasets can take months to years owing to their scale and structural complexity [19]. The situation is more difficult for cryo-FIB-SEM data, as the artifacts and noise can greatly reduce the annotation confidence of human experts. Note that to establish a truly independent validation, manual segmentation needs to be performed on raw cryo-FIB-SEM images, so it is free from any bias potentially introduced by the artifact removal and noise suppression processes. Therefore, we only present the results for visual comparison, and neither detection accuracy nor the correctness of reconstructed membrane topology can be quantitatively established.

## 3 Discussion

In this paper, we present a post-acquisition strategy for restoring cryo-FIB-SEM volumes using structural supervision from comparatively clean, RT-FIB-SEM data. Synthesis guided by experimental artifacts allowed us to construct separate training tasks for artifact suppression and membrane contrast enhancement. The improvements observed on held-out synthetic data, together with the changes in membrane visibility across four acquired cryo-FIB-SEM datasets, suggest that conventional vEM data can provide a useful source of supervision for the restoration of noisy cryo-FIB-SEM volumes.

The distinction between the two stages is important beyond the choice of network architecture. Charging-associated distortions, directional streaks and shading can extend over large fields and depend on their intensity contrast with the surrounding image. Their correction benefits from preserving those relationships rather than removing the background variations in advance. Membrane signal enhancement, by contrast, can emphasize local structural information after broad intensity disturbances have been reduced, and can use adjacent slices as additional evidence. This division complements existing approaches to charging correction and cryo-FIB-SEM processing [5, 6, 8], without requiring modifications to image acquisition. It also provides a way to construct separate synthetic supervision targets for two objectives that would otherwise be difficult to specify jointly.

An important motivation for this work was to make restored cryo-FIB-SEM data more accessible to analysis tools developed for conventional vEM. Using the pretrained CebraNET membrane predictor [12], we observed membrane predictions that more closely followed visible structures in the Stage-2 outputs than in the original inputs (Fig. 5). This is a practical indication that image restoration using our models can facilitate downstream image interpretation, rather than merely changing visual contrast. Potentially, this can also greatly reduce the effort needed to develop various data interpretation tools specifically for cryo-FIB-SEM datasets.

Interpretation of restored cryo-FIB-SEM volumes must remain tied to information present in the acquired images. The quantitative synthetic tests measure agreement with constructed references, including a computationally enhanced membrane target, and should not be read as pixel-level estimates of accuracy on cryo-FIB-SEM datasets. When a region is severely obscured by charging or halos across the central and neighboring slices, the underlying morphology cannot be reliably inferred; introducing a plausible contour would not establish that the contour was present in the specimen. Representative examples contrast persistent information loss across consecutive slices with regions where neighboring slices retain partial structural evidence (Supplementary Fig. 2). Similarly, the 2D overlays and 3D renderings of CebraNET membrane predictions shown here are qualitative evidence of downstream utility, not verification of segmentation accuracy. Examining the original data, intermediate corrections and consecutive restored slices together remains important when interpreting biological structures.

Two technical directions are particularly relevant for extending this workflow. First, axial resampling is presently used only to prepare anisotropically sampled volumes for downstream 3D segmentation, not to reconstruct additional acquired information. In our exploratory analyses, increasing the density of interpolated slices could introduce spurious membrane-like predictions, underscoring the need for interpolation that preserves plausible membrane evolution without propagating isolated noise or creating false continuity along the z-axis. Second, residual local displacement remains a challenge for multi-slice refinement. Our current combination of synthetic drift augmentation and explicit, center-referenced local alignment reduces reliance on perfectly registered inputs, but does not fully resolve spatially varying correspondence. Learning alignment within the refinement model is a promising direction, provided that increased robustness to drift does not come at the expense of membrane sharpness, as encountered in preliminary development. Addressing these trade-offs could further improve the reliability of cryo-FIB-SEM restoration in 3D. A robust z-axis interpolation method might also make it possible to reconstruct the same 3D volume with more sparsely sampled images, potentially reducing the number of slices needed to visualize the same sample and increasing the data collection throughput.

## 4 Methods

### 4.1 Generation of cryo-FIB-SEM Datasets

Four cryo-FIB-SEM volumes were used in our experiments: the publicly archived *Chlamydomonas* dataset EMPIAR-11275 [16], and three in-house datasets of HSAECs, mouse brain tissue and a worm–algae symbiotic system (Fig. 4). The in-house volumes were acquired at a voxel size of 5×5×20 nm. Before analysis, they were binned twofold in X and Y, with no binning in Z, giving voxels of 10 × 10 × 20 nm. EMPIAR-11275 was analysed unbinned (5 × 5 × 20 nm).

#### HSAEC Culture and Vitrification

Primary human small airway epithelial cells (HSAEC, ATCC PCS-301-010) were cultured in airway epithelial cell basal medium (ATCC PCS-300-030) supplemented with the bronchial epithelial cell growth kit (ATCC PCS-300-040) at 37 °C in a humidified atmosphere with 5% CO_2_. EM grids (Quantifoil R 2/2, 200 mesh Au) were glow-discharged (PELCO easiGlow, Ted Pella). Each grid was then placed on a 50-µl droplet of fibronectin (0.2 mg/ml in PBS; Sigma-Aldrich) in a 10 cm dish, incubated for 30 min at 37 °C and 5% CO_2_, and rinsed twice with PBS. The coated grids were transferred to a 35 mm dish containing 2 ml of medium, seeded with 80,000 cells and incubated for 24 h. Grids were back-blotted for 9 or 12 s at 22 °C and 95% relative humidity, plunge-frozen in liquid ethane using a Leica EM GP2, and stored in liquid nitrogen.

#### Mouse Brain Tissue Preparation and Vitrification

All animal procedures were approved by the Stanford University Administrative Panel on Laboratory Animal Care (APLAC; protocol #33824). *AppSAA* knock-in mice were obtained from The Jackson Laboratory (B6.Cg-*App*^tm1.1Dnli^/J, strain #034711; RRID:IMSR JAX:034711). The line carries a humanized Aω sequence plus three familial Alzheimer’s disease mutations knocked into the endogenous mouse *App* gene: Swedish (KM670/671NL), Arctic (E693G) and Austrian (T714I), on a C57BL/6J background [17]. One 12-month-old female homozygous (KI/KI) mouse was used. The mouse was anesthetized with an intraperitoneal injection of 2.5% Avertin (2,2,2-tribromoethanol) and transcardially perfused with cold 0.9% saline for 2 min to remove blood. The brain was then dissected and cut into 100-µm-thick coronal sections in Hibernate-A medium (Gibco, A1247501) at 4 °C using a vibratome (Leica VT1200S). Tissue disks were punched from somatosensory cortex with a 2-mm biopsy punch in a Petri dish. Each disk was transferred to a 200-µL drop of Hibernate-A medium and picked up with a Quantifoil R2/2 grid. The grid was back-blotted with filter paper for 10 s and transferred into a Leica type B carrier (6 mm diameter, 200 µm deep) filled with 2-methylpentane (M32631, Sigma-Aldrich). The carrier was capped with the flat side of a type A carrier, and the sample was high-pressure frozen (Leica EM ICE) at 2,100 bar.

#### Worm–Algae Symbiotic System Vitrification

Worms kept in saline medium were transferred into 6 mm Type B gold-plated specimen carriers that had been lubricated with hexadecene, and then capped with the flat side of a type A carrier. They were vitrified with 5% glycerol, which served as both cryoprotectant and cavity filler. Vitrification was performed similar to the mouse brain tissue, using a Leica EM ICE high-pressure freezer.

#### Cryo-PFIB-SEM Volume Imaging

All in-house datasets were acquired using a Hydra Bio cryo-Plasma FIB-SEM (cryo-PFIB; Thermo Fisher Scientific) at the Stanford-SLAC Cryo-ET Specimen Preparation Center (SCSC). The instrument has a multi-species plasma ion source (Xe, Ar, O, N), an electron column with a magnetic immersion lens, a gas injection system (GIS) and a Pt micro-sputter coater, in addition to a cryostage and SubAngstrom GEN2 large dewar for extended cryogenic operations. Vitrified samples were clipped into AutoGrids, loaded into a 35° pre-tilted shuttle under liquid nitrogen, and transferred to the microscope under cryogenic conditions.

Regions of interest were identified from SEM tile sets acquired with Maps 3.33 (Thermo Fisher Scientific). For mouse brain tissue, the region of interest was transferred to a gold grid by cryo-lift-out as described previously [21]. To protect the specimens, five alternating layers were applied: micro-sputtered Pt, GIS, Pt, GIS, Pt (GIS 2 min per layer; sputtering 4–10 min per layer). This gives a smooth, conductive surface, which reduces curtaining and charging and makes autofocus more reliable.

Xe^+^ plasma (30 kV, 5–10 nA) was used to expose the region of interest and prepare the block face at a 20° milling angle. Serial sectioning used Ar^+^ plasma at 30 kV with a 20 nm step, which produces smooth cross-sections. After each slice, an SEM image was acquired automatically with Auto Slice and View 5 (Thermo Fisher Scientific), using the through-lens detector (TLD) in immersion mode at a 5 nm pixel size. Detector contrast and brightness were kept constant within each acquisition. Dataset-specific parameters are given in Table 1.

**Table 1.** Dataset-specific parameters.

| **Parameter** | **HSAEC** | **Mouse brain** | **Worm–algae** |
|---|---|---|---|
| Vitrification | Plunge freezing | High-pressure freezing | High-pressure freezing |
| Ar<sup>+</sup> milling current | 200 pA | 200 pA | 200 pA |
| SEM voltage | 1.2 kV | 1.2–1.4 kV | 1.2–1.5 kV |
| SEM beam current | 13 pA | 13 pA | 13 pA |
| Dwell time | 100 ns | 50 ns | 50–150 ns |
| Line integration | 128 | 150 | 100–150 |

### 4.2 Data partitioning

We used the publicly available jrc mus-liver-6 mouse-liver volume from OpenOr-ganelle as the source of biological ultrastructure for synthetic supervised training and evaluation [2]. This conventionally prepared, RT-FIB-SEM dataset was used as a structural reference, not as artifact-free cryo-FIB-SEM ground truth. From its finest-resolution, isotropic 8 nm image level, we extracted 22 spatially separated blocks, each consisting of 16 consecutive 1,024 × 1,024 eight-bit slices. Entire blocks were assigned to training (16 blocks), validation (3 blocks), or held-out testing (3 blocks); this partition was shared by both restoration stages. Stage-1 training pairs and Stage-2 five-slice training examples were synthesized independently from their respective source blocks. Held-out test blocks were excluded from model fitting, checkpoint selection, and the training-domain reference statistics used for experimental data inference.

Acquired cryo-FIB-SEM images provided empirical guidance for the artifact models. Charging-artifact characterization was based on a source volume from the worm-algae symbiotic system dataset. All charging cores were manually annotated, yielding binary core masks from which observed core appearances and associated scan-direction intensity residuals were extracted. To separate source templates across experimental partitions, different slices of this volume were used for Stage-1 training, validation, and synthetic testing. Other degradation families, including stripe-like and spatially varying intensity distortions, were generated procedurally using parameterized mathematical models informed by visual observations across experimental cryo-FIB-SEM datasets, rather than by extracting dataset-specific templates. The artifact characterization and synthesis procedures are detailed in Methods 4.3 and 4.4.

### 4.3 Experimental charging-artifact characterization and template extraction

Charging in cryo-FIB-SEM can introduce localized dark regions accompanied by elongated, scan-direction-dependent intensity distortions [5, 7]. To construct the empirical charging-artifact library, we used manually annotated cryo-FIB-SEM slices from a dedicated region of the worm-algae symbiotic system dataset (Methods 4.2). Each annotation was represented by a binary mask identifying the observed core rather than a fitted geometric shape. Eight-connected annotated regions containing at least five pixels were treated as individual cores. We analyzed local background, core intensity, and adjacent dark and bright intensity deviations to characterize their spatial extent and relative contrast. Bright positive deviations were recorded as halo statistics but were not extracted as a separate real-valued template; the stored library consisted of observed core appearances and associated dark-tail residuals.

Nearby annotated cores were grouped before template extraction to preserve multiple charging centers and potentially overlapping scan-direction tails. For each core, a grouping rectangle was centered on its annotation centroid, with height equal to the core bounding-box height and width equal to the larger of the core width and six times its height. Cores whose grouping rectangles overlapped, directly or through a chain of overlaps, were assigned to the same group. The union of their grouping rectangles was expanded by 12 pixels vertically and 24 pixels horizontally to form an extraction crop. The grouping dimensions were empirically selected for this study and served only to define the extraction region; the core masks remained the exact manual annotations.

For a grouped crop I*_g_*, let M*_g_* denote the union of its annotated core pixels. We preserved the recorded pixel intensities inside M*_g_* as the core template, without smoothing or estimating replacement intensities. To estimate the surrounding dark tail, we constructed a directional reference image I*_g_* by applying horizontal Gaussian filtering to the normalized crop at kernel lengths of 151 and 31 pixels. The mean-centered low-frequency and intermediate-frequency components were subtracted with coefficients 1.0 and 0.75, respectively, and the corrected image was mapped back to the original intensity range. This reference was used only to estimate local intensity deviations; it was not treated as a ground-truth artifact-free image. The preliminary outside-core dark residual was [*Î_g_* − I*_g_*]_+_, with [*a*]_+_ = max(a, 0). We retained only eight-connected residual regions linked to the six-pixel-dilated core neighborhood, using a permissive 0.5-gray-level connectivity threshold, and removed remaining values below 1.0 gray level. The exported tail template was therefore

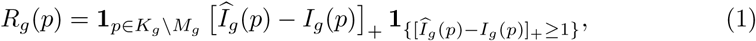

where K*_g_*is the retained connected support and p denotes a pixel location. Core pixels were always excluded from R*_g_*, ensuring that core and tail intensities could be transformed separately during subsequent synthesis.

To preserve the empirical contrast relationship between these components, we measured the local median background intensity b*_g_*, median core intensity c*_g_*, and 90th percentile t*_g_* of positive tail residuals above 1 gray level. The per-template tail-to-core ratio was defined, when the core-to-background contrast was positive, as

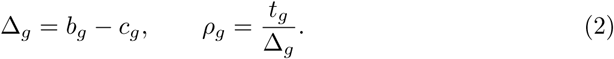

These measurements were stored as metadata rather than used to alter the extracted pixel values. Library entries were retained only when the core image matched the original recorded core pixels, the tail was finite and nonnegative, and at least five tail pixels remained with a total residual of at least 1 gray-level pixel sum. Each accepted entry included the raw crop, original core intensities, binary core mask, dark-tail residual, and associated geometric and intensity statistics. No normalization or spatial resizing was applied to the stored templates. For subsequent Stage-1 synthesis, the library was partitioned by source slice into 461 training entries (slices 0–49), 34 validation entries (slices 50–54), and 39 test entries (slices 55–59) (Methods 4.4).

### 4.4 Stage-1 synthetic training data generation

We constructed paired 2D examples in which biological ultrastructure was provided by the publicly available, conventionally prepared JRC mouse-liver FIB-SEM dataset [2], while imaging artifacts were synthesized separately. The structural data comprised 22 spatially separated blocks of 16 consecutive 1,024 × 1,024 slices, partitioned by block into training (16), validation (3), and testing (3). Because the clean source and experimental cryo-FIB-SEM data differed in intensity distribution, we first adapted each source slice to one of four nominal background medians, L ∈ *{*115, 145, 175, 200*}*. Contrast was estimated from the 5th–95th percentile range of experimental cryo-FIB-SEM reference images, excluding annotated charging cores, and applied as a weak, bounded affine adjustment:

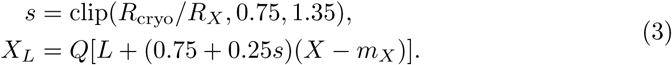

where X is the original clean slice, m*_X_* its median, R*_X_* and R_cryo_ the respective 5th–95th percentile ranges (each floored at 20 gray levels), and Q denotes rounding and clipping to the eight-bit range. Four slices per 16-slice block were assigned to each background level. The transform modified the image intensity statistics without globally sharpening or replacing biological structures.

Charging artifacts were simulated using experimentally derived core-intensity images, binary core masks, dark-tail residuals, and per-template contrast measurements extracted as described in Methods 4.3. For each charging event, a paired core and tail was jointly resized and positioned on the styled slice; the tail was additionally elongated in the horizontal scan direction while the core geometry was retained. A fixed template-resizing factor of 5/8, a morphological multiplier of 2.2, and an additional random scale factor sampled from [0.7273, 1.2] introduced variations in artifact extent; scan-direction tails were independently stretched with proximal factors sampled from [4, 8]. The observed core appearance was placed within its mask after a small contrast adjustment, with its median intensity calibrated to the range 35–70 gray levels. Rather than independently sampling the tail strength, we coupled it to the realized core-to-background contrast, defined as Δ*_c_* = median*_p_*_×*M*_ [X*_L_*(p) −C^↔^(p)], where M is the placed core mask and C^↔^ is the calibrated core-intensity image. Using the measured tail-to-core contrast ratio ε, constrained by empirical reference statistics and clipped to [0.30, 0.70], we set the target tail darkening to

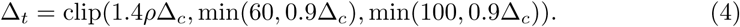

The positive tail residual was then rescaled so that its 90th-percentile darkening matched Δ*_t_*. To simulate both attenuation and partial loss of local contrast, we rendered the non-core charging field using

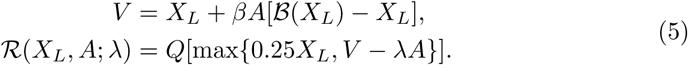

where A is a smoothed, normalized residual-derived opacity map, *B* combines Fourier-domain low-pass filtering with spatial Gaussian smoothing, ω ∈ [0.08, 0.15] controls local structural attenuation, and ϑ is fitted to the target tail darkening. The calibrated raw core pixels were then inserted into the rendered field. Overlapping dark fields were fused by a smooth maximum of background-relative darkening (temperature 4 gray levels) rather than by overwriting earlier artifacts.

We additionally synthesized several artifact morphologies that could co-occur with charging. *Broad shading* was produced from a jointly enlarged core–tail field (an additional 2–3-fold enlargement), without pasting an opaque core. Defining its calibrated dark residual as 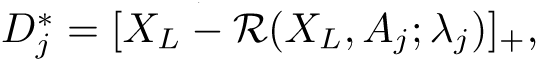 we applied

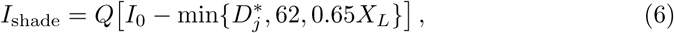

where I_0_ denotes the image before this perturbation, A*_j_*is the combined core–tail opacity field, and the 90th-percentile darkening was calibrated to 50–58 gray levels.

*Thin dark-line bundles* were assembled from approximately horizontal, 1–2-pixel-wide strokes with randomized lengths, spacing, and endpoints. Their softened support A*_b_* included a spatial envelope that weakened surrounding traces (relative strength 0.10–0.26). Unlike broad charging tails, these bundles did not undergo additional structure filtering:

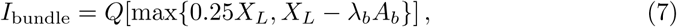

with *λ_b_* chosen for a 90th-percentile darkening of 60–100 gray levels.

*Bright halos* were generated around the same charging core–tail support rather than sampled from an independent halo-template library. The support was anisotropically Gaussian-smoothed, masked to protect existing dark core/tail pixels, normalized, and adjusted by a power transform to obtain H ∈ [0, 1]. Brightening was then applied as

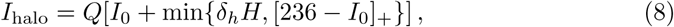

where δ*_h_* ∈ [45, 62] gray levels and the halo-map exponent was sampled from [0.55, 0.72]. Two *mixed-polarity stripe* subtypes were also synthesized. Broad stripes consisted of a bright horizontal seam bordered by wider, weaker dark flanks:

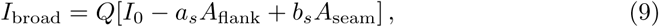

where a*_s_* ∈ [6, 10] and b*_s_* ∈ [8, 18] gray levels. Narrow stripes comprised 4–7 alternating dark and bright bands with variable extent:

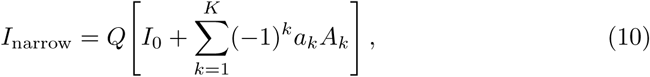

where K ∈ *{*4, 5, 6, 7*}*, A*_k_*denotes a softened band envelope, and a*_k_* > 0 its magnitude. The broad and narrow subtypes were selected with probabilities 0.55 and 0.45, respectively. These stripe perturbations affected only the degraded inputs, not the supervision targets.

The supervised target retained the styled clean image outside the charging-core masks. Within each synthetic charging-core mask, we replaced the underlying image content with background texture selected from the lowest-structure patch among 64 randomly sampled candidates from the same clean slice, together with a 4-pixel-wide dark cue along the inner core boundary. The texture intensity was matched to the surrounding eight-pixel ring, while the boundary intensity was set to m_ring_ − 0.68d*_m_*, where m_ring_ is the local background median. To estimate d*_m_*, we calculated positive local dark contrast relative to a Gaussian-smoothed structure-enhanced reference (σ = 3 pixels), selected pixels where the structure-enhanced reference differed from the styled clean image by at most −2 gray levels, and multiplied the 75th percentile of the selected contrast values by 1.25. The resulting value was clipped to [10, 36] gray levels. This design preserves a simplified membrane-like cue while encouraging restoration to the surrounding background intensity, without assuming that the synthetic boundary represents the actual occluded ultrastructure.

For each nonidentity 1,024 × 1,024 example, the generator placed seven separate charging events based on experimental templates and two dark-line bundles, applied the halos associated with the charging events, then added two broad joint-shading events and three mixed-polarity stripe objects. Negative residuals outside the pasted cores were additionally amplified with background-dependent factors (*{*1.00, 1.15, 1.30*}* for L = 115, 145; *{*1.00, 1.30, 1.50*}* for L = 175; and *{*1.00, 1.35, 1.70*}* for L = 200), without amplifying the original core or positive halos. Four seeded variants were generated per styled source slice, including approximately 15% strict identity pairs (I = Y = X*_L_*) without synthetic artifacts. The resulting datasets comprised 1,024 training pairs (872 degraded, 152 identity), 192 validation pairs (164 degraded, 28 identity), and 192 test pairs (164 degraded, 28 identity). Clean biological source blocks and experimental charging-template source slices were partitioned independently across training, validation, and testing to prevent source reuse between splits.

### 4.5 Stage-1 network architecture and training

Stage-1 was trained as a supervised 2D model for artifact suppression, processing each slice independently without requiring cross-slice registration. We adopted a modified nonlinear activation-free network (NAFNet) [13]. To accommodate complete 1,024 × 1,024 images, a factor-two pixel-unshuffe operation rearranged each single-channel image into four channels at half the lateral resolution; its inverse, pixel shuffe, restored the original pixel arrangement after network processing. These operations rearrange pixels without interpolation. Writing *P*_2_ for pixel unshuffe and R*_ω_* for NAFNet’s learned packed-domain residual, the Stage-1 prediction was

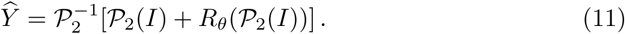

Input and target images were scaled from eight-bit intensities to [0, 1] before training. The network used a base width of 16 channels, four encoder levels containing (1, 1, 1, 2) NAF blocks, a four-block bottleneck, and four decoder levels containing one block each. Encoder–decoder skip connections were added at corresponding resolutions. The NAF blocks retained depthwise convolutions, multiplicative gating and simplified channel attention [13], with dropout probability 0.1 in their two residual branches. The model had 2,658,004 trainable parameters.

We optimized three complementary terms defined using the target correction Ĉ = Ŷ − I and predicted correction C = Y − I:

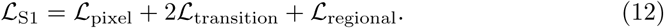

The *difference-adaptive pixel loss* gave greater weight to locations requiring substantial artifact correction. For each pixel p, we defined d(p)= |I(p)−Y (p)|, a(p)= clip[(d(p)− 8/255)/(32/255), 0, 1], and w(p) = 1 + 4(G_2_ ↔ a)(p), where G*_ε_* denotes normalized Gaussian smoothing with standard deviation σ pixels. The weight-normalized loss was

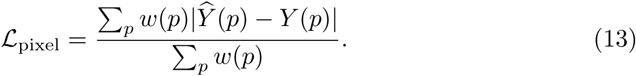

The weight map was fixed by the input–target pair and was not learned.

The *transition-local correction-gradient loss* constrained interfaces between regions requiring corrections of opposite signs, such as the boundaries between dark charging and adjacent bright distortions. We identified pixels with C ≥ 20/255 and C ≤ −20/255; the intersection of their binary dilations (elliptical radius 5 pixels) defined a transition mask T. For each valid horizontal or vertical neighbor pair (p, p + e*_d_*), we set T*_d_*(p) = max*{*T (p),T (p + e*_d_*)*}* and ∇*_d_*C(p)= C(p + e*_d_*) − C(p), for d ∈ *{*x, y*}*. The signed-correction gradient loss was

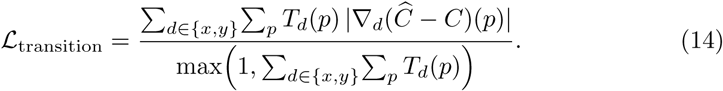

The loss therefore focused on regions near changes in correction sign, rather than applying the gradient penalty across the entire image. The transition masks were computed from the input–target corrections and remained unchanged during training.

The *multi-scale regional correction loss* addressed broad, low-amplitude residual shading. At Gaussian scales σ ∈ *{*16, 32*}* pixels, we selected locations for which 1/255 ≤ |(G*_ε_* * C)(p)| ≤ 8/255 and |C(p)| < 20/255, defining binary masks M*_ε_*. Both positive and negative regional corrections were included, while strongly corrupted pixels were excluded. We compared smoothed correction fields using

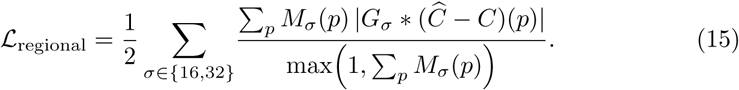

All losses were computed over the entire slice, with no clipping of predicted intensities before loss evaluation. The pixel, transition and regional terms respectively emphasized overall fidelity, local changes in correction sign, and weak spatially extended intensity corrections.

Training used 1,024 full-resolution paired images (872 degraded and 152 identity pairs); the separate validation and test partitions each contained 192 pairs (164 degraded and 28 identity), as described in Methods 4.4. We used AdamW [22] with initial learning rate 10^−4^ and weight decay 10^−4^. Each microbatch contained one full-resolution 1,024 × 1,024 training image, and gradients were accumulated over four microbatches (effective batch size four), with global gradient-norm clipping at 1.0. Stage-1 training was performed on an Apple Silicon GPU using the Metal Performance Shaders (MPS) backend. Training used random seed 123 and a maximum of 200 epochs. The learning rate was halved on a validation-loss plateau (patience six epochs, relative threshold 0.003), down to 6.25 × 10^−6^. Early stopping was permitted after at least 25 epochs, two learning-rate reductions and ten epochs without a meaningful validation-loss improvement (relative threshold 0.003). Training stopped after 148 epochs, and the checkpoint from epoch 146, which achieved the *lowest total validation loss*, was retained for inference. PSNR was monitored but was not used for model selection.

### 4.6 Stage-2 reference construction and synthetic degradations

Stage-2 training used the same JRC mouse-liver FIB-SEM source and block-level data partition as Stage-1 [2], but its inputs and supervision targets were generated independently of Stage-1 predictions. Each training example consisted of five 256 × 256 patches cropped from consecutive source slices, R*_z_*_−2_,…,R*_z_*_+2_, with the central slice R*_z_* providing the reference for target construction. Before patch extraction, each complete raw slice was standardized using its own mean µ*_j_* and standard deviation s*_j_*, followed by Gaussian low-frequency subtraction and histogram-mode centering. With N*_j_* = (R*_j_* − µ*_j_*)/s*_j_* and m*_j_* = mode_50_(N*_j_*− G_10_ * N*_j_*), the processed slice was

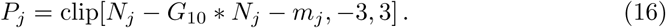

Here G*_σ_* is Gaussian smoothing with standard deviation *σ* pixels, * denotes convolution, and mode_50_ is the midpoint of the most populated bin in a 50-bin histogram. These statistics were computed on entire 1,024 × 1,024 slices, not separately on individual 256 × 256 training patches. The same full-slice intensity preprocessing was used for Stage-2 experimental data inference (Methods 4.8).

The supervision target was a deterministic, structure-enhanced version of the *original clean central slice*, constructed independently of the sampled input degradation. A bilateral filter [23] (diameter 7 pixels; intensity/spatial standard deviations 12 gray levels/2 pixels) yielded B*_z_*, from which signed local detail was extracted as D*_z_* = B*_z_* − G_2_ * B*_z_*. We defined the smoothed enhancement gate q*_z_* = G_0.8_ * h(clip[(|D*_z_|* − 1.5)/2.5, 0, 1]), where h(t)= 3t^2^ − 2t^3^. The enhanced eight-bit reference was

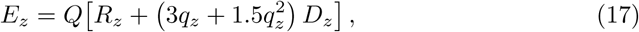

where Q rounds and clips intensities to [0, 255]. The training target was expressed in the *unenhanced* center slice’s normalization basis, rather than normalized again from E*_z_*:

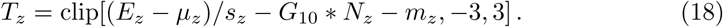

This reference enhances locally prominent image detail without segmentation annotations. It is a computationally enhanced RT-FIB-SEM supervision target, not a measurement of ultrastructure obscured by artifacts in experimental cryo-FIB-SEM images.

For missing-region synthesis, crop locations and occlusion centers were preferentially selected from regions with pronounced dark, high-gradient structures. An annotation-free placement score combined raw-image darkness, negative processed intensities and Sobel gradients, using the center slice (weight 0.72) and its four neighbors (combined weight 0.28). Of 24 proposed crops, one was sampled from the six highest-scoring candidates with probability proportional to its score; within the selected crop, occlusion centers favored pixels above the 78th percentile of the structural score. Conditional on sampling a missing-region example, four patch-level patterns were used: center only (50%), center dominant with weaker neighboring corruption (25%), neighbors only (15%), or irregular multi-patch corruption (10%). Affected patches received individually perturbed, irregular masks with nominal base area 2,500–8,000 pixels^2^, scaled by a pattern-dependent severity and softened with a 2.5-pixel Gaussian. Their replacement fields combined the median of a surrounding ring (approximately 6–18 pixels beyond the mask), a small intensity offset and weak low-frequency random texture. This simulated local information loss rather than a uniform black or zero-valued cutout. The soft-mask blending operation is specified below because it occurs after Gaussian noise is added.

Two additional principal degradations modeled inter-slice inconsistency. For *local drift*, the central crop and target remained fixed, while one to four neighboring crops were sampled from spatially shifted positions in their original preprocessed full slices. For neighbor j, with an integer-rounded translation **δ***_j_* of randomly oriented magnitude 4–15 pixels, the shifted patch over crop region Ω was

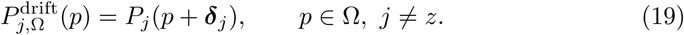

Valid crop bounds were respected; no image warping or subpixel interpolation was performed. For *loss of focus*, exactly one slice was selected, with probability 0.50 for the center, 0.15 for each immediate neighbor and 0.10 for each outer neighbor. Blurring was applied to the *complete raw slice* before repeating the full-slice preprocessing and extracting the patch:

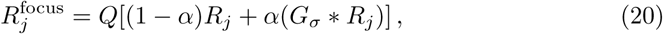

with *σ* ∼ *U*(4.0, 5.8) pixels and *α* ∼ *U*(0.92, 0.96). The clean enhanced target T*_z_* was unchanged even when the central input was blurred.

Residual scan-direction artifacts were synthesized independently of the principal condition. Let *P_j_*’ denote the processed input patch after applying the selected principal condition, before adding residual stripes or noise. A categorical draw selected no composite stripe (30%), original polarity with a bright center and dark flanks (35%), or inverted polarity with a dark center and bright flanks (35%). For each input assigned a stripe polarity, we sampled K ∈ *{*4,…,7*}* approximately horizontal stripe objects. Their contributions were accumulated and clipped after all K objects:

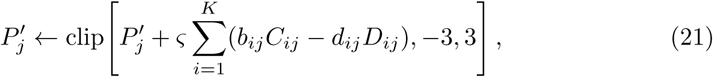

where φ = +1 or −1 sets the original or inverted polarity, respectively, C*_ij_* and D*_ij_* are soft core and flank masks, and b*_ij_* and d*_ij_* their sampled strengths. Stripe geometry and amplitude varied slightly across the five input patches; opposite polarities were never combined within the same five-patch training example. An additional sparse thin-line bundle was independently introduced with probability 0.15, using discontinuous horizontal strokes 1–2 pixels thick and 80–190 pixels long. Its high-pass-like intensity signature, proportional to −V +G_10_ *V for a synthetic dark-line field V, was added at low strength (nominal darkening 0.32–0.62 processed-intensity units). *Gaussian noise* was a separate degradation present in *every* training input, including examples with no principal structural corruption:

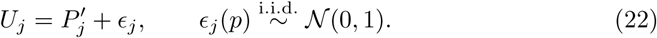

Noise was sampled independently across pixels and the five input patches in processed-intensity units.

For each five-patch training example, exactly one principal condition was drawn: no principal structural corruption (40%), missing region (30%), local drift (15%) or loss of focus (15%). Composite stripes and thin-line bundles were then independently sampled as described above, followed by Gaussian noise. Only for missing-region examples, the precomputed soft masks M*_j_* ∈ [0, 1] and locally matched fill fields F*_j_* were applied *after* noise injection:

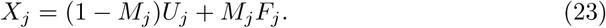

For other examples, X*_j_* = U*_j_*. Thus, noise remained outside missing regions but was largely replaced inside their fully obscured portions. Each degraded five-channel input **X** ∈ R^5×256×256^ was paired with the fixed enhanced central target T*_z_*|_Ω_. The network and optimization procedure are described in Methods 4.7.

### 4.7 Stage-2 network architecture and training

Stage-2 used a five-channel, 2D U-Net-like encoder–decoder [14] to predict one membrane-enhanced central patch from a five-slice patch input. Given the syn-thetically degraded, preprocessed patches described in Methods 4.6, its prediction was

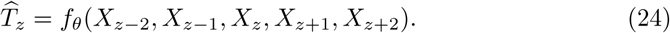

The network performed early fusion of all five input patches through its first convolution, rather than using separate encoders or 3D convolutions. Four encoder levels had 32, 64, 128 and 256 channels, with 2 × 2 max-pooling between resolutions and a 256-channel bottleneck. The decoder used four transposed-convolution upsampling stages and multiscale feature concatenation. Convolution kernels were 5 × 5 at the first three encoder levels and output head, and 3 × 3 at the deepest encoder level and bottleneck; decoder upsampling used 3 × 3 or 5 × 5 kernels. Hidden activations used the sine function, with elementwise dropout of 0.1 [24]. The network contained 2,204,289 trainable parameters and was initialized from scratch using truncated LeCun-normal weights and zero convolutional biases [25].

To retain a direct full-resolution reference from the central input patch, the network used a skip connection at the final decoder stage. The final 32-channel upsampled decoder representation was concatenated with *only* the processed central input patch,

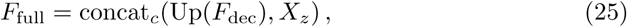

where Up denotes the last learned upsampling layer and concat*_c_* channel concatenation. A convolutional prediction head mapped the resulting 33-channel representation to one output channel. Neighboring patches contributed through fused encoder features and intermediate skips, while this direct pathway preserved the central patch as the full-resolution spatial reference. Importantly, the connection was concatenative: the network predicted a complete intensity image, without adding the central input as an output residual or applying a final sigmoid.

Training used 256 × 256 target patches cropped from the structure-enhanced central-slice reference T*_z_* defined in Eq. (18). For predictions T̂ and targets T, we minimized

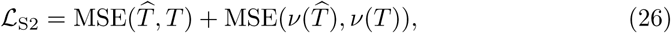

where *v*(u) = max(−u, 0) and each mean squared error is averaged over all pixels and examples in the batch. The first term measured prediction error against the enhanced reference; the second additionally penalized differences between the negative-valued components of the prediction and target in the processed intensity domain. This extra term was not a membrane-label-based loss. A separate missing-region masked loss used the same two error terms within binary central-patch masks obtained by thresholding the synthetic missing-region soft masks at 0.5. This loss was recorded for diagnostics only and did not contribute to backpropagation or checkpoint selection.

We trained using Adam [26] (learning rate 2 × 10^−5^, *β*_1_ = 0.9, *β*_2_ = 0.999, *∈* = 10^−8^, zero weight decay), batch size 32 and 256 × 256 patches. The training schedule comprised 2,000 optimizer steps, or 64,000 stochastic patch exposures rather than 64,000 distinct biological patches. Random seed 0 was used, with separate seeded streams for stripe and thin-bundle synthesis. Training used float32 arithmetic, without a learning-rate scheduler, early stopping or Stage-1 weight initialization. As in Stage-1, Stage-2 training was performed on an Apple Silicon GPU using the Metal Performance Shaders (MPS) backend. The fixed validation set consisted of 32 sampled locations from the three held-out validation blocks, each evaluated under noise-only, missing-region, local-drift and defocus conditions; additional original-stripe, inverted-stripe and thin-bundle examples were retained for visual diagnostics. Validation was performed every 100 optimizer steps with dropout disabled. If V*_k_* denotes the mean full-patch loss from Eq. (26) under condition k, the checkpoint-selection score was

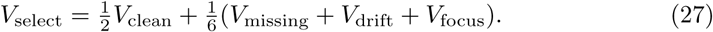

The checkpoint from step 1,900 achieved the lowest validation-selection score and was retained as best.pt. Stripe diagnostics and the missing-region masked metric were excluded from this selection score; held-out test blocks were excluded from both optimization and checkpoint selection.

### 4.8 Experimental data preprocessing and two-stage inference

The four experimental cryo-FIB-SEM datasets evaluated in this study were processed using the trained Stage-1 and Stage-2 checkpoints without updating either network’s weights. For these datasets, source images were converted to eight-bit grayscale TIFF stacks in ZY X order using a single intensity window estimated from the 0.5th–99.5th percentiles of pixels sampled across the entire volume, rather than independent per-slice rescaling. Each converted volume was then processed using variational stationary noise removal (VSNR) [15] to suppress curtaining before learned restoration. The VSNR implementation used a Gabor filter with transverse and longitudinal scales of 1 and 30 pixels, noise level 80 and up to 30 iterations, with optional orientation selection over −5^≃^ to 5^≃^; corrected slices were robustly matched back to their original median and median absolute deviation. Unusable slices could be excluded and a region of interest selected before inference.

The three in-house cryo-FIB-SEM datasets underwent global translation-only alignment before Stage-1 to reduce lateral displacement between successive slices. Pairwise XY translations were estimated between nearby slices using multiscale phase correlation [27] and combined to obtain a global translation **t***_z_* = (t*_x,z_*, t*_y,z_*) for each slice. The aligned image was obtained as

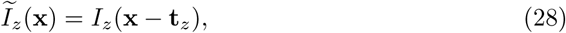

where **x** denotes the in-plane pixel coordinates. Translations were applied using bilinear interpolation, followed by cropping to the common valid field of view where needed. No rotation or nonrigid registration was performed. The publicly available EMPIAR-11275 volume was already spatially aligned and therefore did not undergo additional global alignment.

Global alignment is an optional preprocessing step, and users may choose an alternative alignment method according to their dataset or omit this step for data that are already aligned. This stack-level operation is distinct from the patch-local center-referenced alignment used during Stage-2 inference.

*Stage-1 artifact suppression* operated independently on each aligned or otherwise prepared slice, as described in Methods 4.5. Each eight-bit slice was divided by 255 and passed to the trained 2D model without additional intensity-style adaptation. Slices with both dimensions at most 2,048 pixels were processed at native resolution. Larger slices were processed in overlapping 2,048 × 2,048 tiles (512-pixel overlap; 1,536-pixel stride). To avoid seams, the model-predicted correction Δ*_k_*= f*_ω_*_1_ (X*_k_*) −X*_k_*from each normalized input tile X*_k_* was combined using normalized Hann weights w*_k_*:

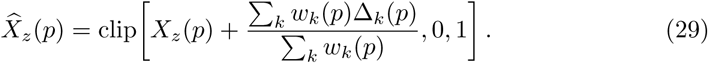

Only tiles covering pixel p contribute to these sums; the blending weights were floored at 10^−3^ near tile boundaries. The restored slices were converted back to eight-bit intensities to form the Stage-1 output volume.

*Stage-2 intensity preprocessing* was performed on each complete Stage-1 output slice before extracting any inference patches. The operation was identical to the full-slice normalization, Gaussian low-frequency subtraction (σ = 10 pixels), 50-bin histogram-mode centering and [−3, 3] clipping used for Stage-2 training (Eq. (16)). For the experimental datasets reported here, we enabled an optional intensity-and Fourier-statistics matching step to match the processed inputs to the clean training-reference statistics. This procedure comprised two ordered operations: Fourier log-amplitude distribution matching followed by spatial-intensity distribution matching. Although related to Fourier-based domain adaptation [28], this method matched global empirical distributions rather than replacing an image’s low-frequency Fourier coefficients with those of a single reference image.

Both mappings used the clean JRC liver *training* split as their target domain, without using validation or test blocks. We sampled 128 clean training slices (seed 20260924), preprocessed each complete slice and extracted one random 256 × 256 reference patch per slice. Histograms with 4,096 bins characterized their processed intensities and non-DC Fourier log-amplitudes. For the source domain, Fourier statistics were accumulated over all overlapping 256 × 256 tiles (stride 128) of every preprocessed slice in the current Stage-1 volume. Let F*_k_* be one tile’s Fourier coefficient, Φ*_k_*= arg F*_k_*, and *Q_F_* = (C_t_^F^)^-1^ ° C*_s_*^F^ the empirical source-to-target log-amplitude quantile mapping. For nonzero frequencies, we applied

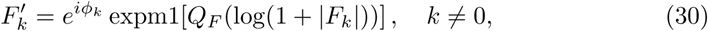

while retaining the original DC coefficient. Inverse transforms of the tiles were assembled by overlapping Hann-weighted blending to give a full-slice result P*_z_*^F^. A second source CDF *C_s_^I^* was measured across *all pixels of all post-Fourier slices* and mapped to the clean-reference intensity CDF *C_t_^I^*:

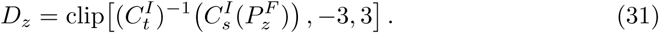

The resulting maps were fixed during inference and did not update either network. Intensity-and Fourier-statistics matching was enabled for the reported experimental-data workflow. Representative comparisons of Stage-2 outputs with and without statistics matching are shown in Supplementary Fig. 3. Because the source CDFs were estimated from the entire supplied Stage-1 volume, changing the selected slices or spatial crop can change the mapping and subsequent predictions.

*Patch-local center-referenced alignment* was applied when assembling each five-slice Stage-2 input. At a given 256 × 256 output-tile location, the central patch remained fixed, and the crop coordinates of the four neighboring patches (offsets *±*1, *±*2) were adjusted independently relative to it. Matching used normalized cross-correlation (NCC) [29] on a central 160 × 160 region within a 320 × 320 context, after Gaussian smoothing (σ = 1.25 pixels) for displacement estimation. For each neighbor, the candidate integer translation was

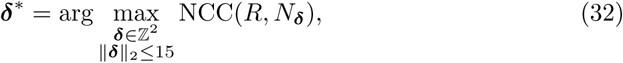

with candidates restricted to those leaving the final neighbor crop within the actual image bounds. A nonzero displacement was accepted only if its NCC was at least 0.15 and improved on zero displacement by at least 0.020; otherwise the neighbor remained unshifted. Accepted shifts altered only the *integer coordinates* used to sample neighboring patches, without interpolation or movement of the center patch. At volume boundaries, missing neighboring context positions were filled by replicating the nearest acquired slice. This patch-local operation compensated for residual lateral differences without generating intermediate slices or performing nonrigid reconstruction.

Finally, the trained five-channel Stage-2 network (Methods 4.7) predicted one central patch from each locally aligned input. Predictions T*_z,k_* from overlapping 256 × 256 tiles (128-pixel stride) were merged in the fixed center-slice coordinate system:

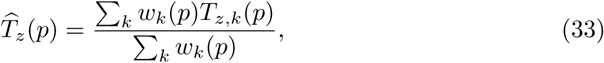

using Hann weights floored at 10^−3^, including terminal tiles along image edges. The processed output was converted to eight-bit intensities by a fixed, dataset-independent decoding rule,

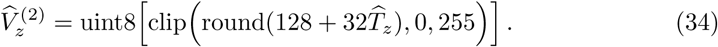

The original low-frequency intensity field was not added back after decoding. Consequently, the final Stage-2 image represented an enhanced, canonically scaled output rather than the original detector-intensity distribution. Stage-1 and Stage-2 outputs were saved separately for comparison. Both stages retained the acquired slices’ ordering and axial sampling; any subsequent z-axis resampling for segmentation was performed independently of the two-stage processing.

### 4.9 Evaluation protocols and quantitative metrics

Both restoration stages were evaluated using the three JRC mouse-liver blocks reserved for testing, which were excluded from model fitting and checkpoint selection. These blocks were separate from the 16 training and three validation blocks. We evaluated each stage against its own synthetic supervision target, rather than against paired, artifact-free cryo-FIB-SEM acquisitions.

For *Stage-1*, we evaluated the fixed checkpoint on all 192 paired 1,024 × 1,024 images in the held-out synthetic test split described in Methods 4.4. Of these, 164 contained synthetic, potentially overlapping artifacts and were pooled for the primary quantitative analysis; results were not stratified by artifact type. We applied the same native-size, 2D inference used for experimental data and calculated metrics from the final clipped and rounded eight-bit predictions against the corresponding synthetic targets. The remaining 28 artifact-free identity pairs were evaluated separately as a preservation control. They were excluded from the primary PSNR summary because their input images exactly equaled their targets, giving infinite input PSNR.

For *Stage-2*, each of the three held-out 16-slice blocks provided 12 valid center positions (z = 2,…,13), with four 256×256 crops per position (two structure-enriched and two spatially random), yielding 144 base patches. Each base patch was tested under six *separate* conditions: Gaussian noise alone, missing region with noise, defocus with noise, local drift with noise, original-polarity stripes with noise, and inverted-polarity stripes with noise. The same original five-slice source context, enhanced central target patch, and Gaussian-noise realization were used for the six matched conditions of each base patch. Synthetic degradation distributions followed Methods 4.6; conditions were not combined within this test. The optional intensity matching and patch-local alignment used during Stage-2 inference were not applied in this controlled synthetic evaluation. Five of these conditions are presented in the main figure (Fig. 3); local drift was also evaluated but is not shown. Degraded central input patches, model outputs, and membrane-enhanced central target patches were all converted to the canonical eight-bit intensity representation Q(128 + 32u) before metric calculation, where Q rounds and clips to [0, 255].

For each image or central patch, we measured mean absolute error (MAE) in gray levels, peak signal-to-noise ratio (PSNR), and single-scale structural similarity (SSIM) [30]. PSNR was calculated as 10 log_10_(255^2^/MSE) in decibels. Both stages used a data range of 255, but their archived SSIM implementations differed: Stage-1 used an 11 × 11 Gaussian window (σ = 1.5 pixels), with the valid central region averaged, whereas Stage-2 used the default seven-pixel uniform window of scikit-image. In addition to full-patch metrics, missing-region performance was quantified within the *central* binary occlusion mask M:

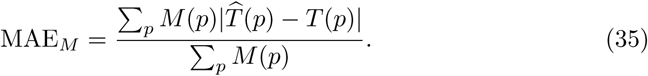

The masked-MAE analysis included only the 118 missing-region test patches with nonempty central masks. The separate center-defocus SSIM analysis included only the 71 test patches in which the central source slice, rather than a neighboring slice, was blurred. All 144 patches were retained in the corresponding full-condition summaries.

Metrics were calculated for each image or patch before aggregation. We report arithmetic means and sample standard deviations (n−1 denominator); gains and error reductions were computed as paired differences for each example before averaging. No inferential significance tests were performed. Multiple patches from the same biological block are correlated and were not treated as independent biological replicates. Because the acquired cryo-FIB-SEM datasets had no paired artifact-free ground truth, their visual comparisons and downstream membrane predictions were not assigned synthetic-reference reconstruction accuracy metrics.

### 4.10 Downstream membrane segmentation and volumetric visualization

We assessed the downstream utility of image restoration with the publicly available pretrained CebraNET membrane predictor from the CebraEM workflow [12, 20], using the weights.torchscript checkpoint archived on Zenodo (DOI: https://doi.org/10.5281/zenodo.10047209). The supplied TorchScript weights were held fixed throughout; we neither retrained nor fine-tuned the segmentation model. For each experimental dataset, corresponding original and Stage-2-restored eight-bit volumes were processed using identical input geometry, interpolation settings, model weights, prediction aggregation and threshold. This comparison evaluated the use of an existing membrane predictor on the restored images, rather than a new segmentation method.

To reduce axial sampling anisotropy for CebraNET inference, we constructed an endpoint-aligned Z grid while retaining the analyzed in-plane pixel spacing. We empirically selected an axial scaling factor of 1.5 for both experimental datasets used in the downstream segmentation analysis. These settings did not produce isotropic voxel sampling. Intermediate planes were synthesized by motion-aware interpolation between consecutive acquired slices. For each pair, the two slices were standardized independently by their means and standard deviations and Gaussian-smoothed (σ = 1.25 pixels) for matching. Bidirectional displacement fields **f***_z→z_*_+1_ and **f***_z_*_+1→*z*_ were then estimated with normalized cross-correlation on overlapping 160×160-pixel patches (80-pixel stride; maximum integer displacement *±*15 pixels per axis). Matches required an NCC score of at least 0.15 and an improvement of at least 0.01 over zero displacement; rejected estimates were filled from nearby accepted matches, and the full-resolution displacement fields were Gaussian-smoothed (σ = 12 pixels). At a fractional slice position z + t, 0 < t < 1, we generated

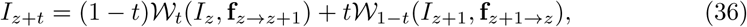

where *W_a_*(I, **f**)(**x**) = I(**x** − a**f** (**x**)) denotes bilinear backward sampling. The first and last planes coincided with the original endpoints. Displacement-confidence and forward–backward consistency measurements served as diagnostics only; they did not weight the synthesized intensities. The interpolated planes were rounded and clipped to the eight-bit intensity range before segmentation inference. These intermediate planes were used solely to prepare the segmentation-model input and were not interpreted as experimentally observed ultrastructure.

Each resampled volume was divided by 255 and processed with the fixed CebraNET model using overlapping 64 × 64 × 64-voxel patches (stride 16, nominal batch size 8). Model inference used CUDA half-precision autocasting; overlapping predictions were accumulated in single precision using separable triangular spatial weights, with a per-axis edge-weight floor of 0.20. The resulting prediction map was then resampled *back* to the original acquired slice positions by linear interpolation of probabilities, before thresholding. Specifically,

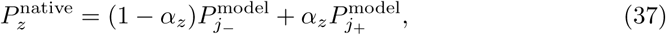

where j_−_ and j_+_ bracket native slice z on the model grid and ς*_z_*is its fractional position between them (or zero when the positions coincide). The native-grid binary membrane mask was defined by 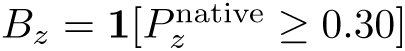 and saved with the same ZY X dimensions as the input volume. No interpolated Z planes were retained in the final segmentation masks.

For 2D inspection (Fig. 5), native-grid binary predictions were displayed as cyan overlays on the corresponding original or restored slices. For volumetric visualization (Fig. 5), predicted masks from CebraNET were converted to smoothed scalar volumes and rendered as surfaces in UCSF ChimeraX [31]. Smoothing was applied for display only, after segmentation; it did not alter the saved probability maps or binary masks. Algal cells in the worm-algae symbiosis dataset and mitochondria in the mouse brain dataset were annotated manually in UCSF ChimeraX using the Volume Eraser tool. Because no manual membrane ground truth was available for these experimental cryo-FIB-SEM volumes, the overlays and surfaces were interpreted as qualitative predictions rather than direct measures of segmentation accuracy or verified membrane topology.

## Data availability

The *Chlamydomonas* dataset is publicly available from EMPIAR-11275. The in-house datasets of mouse brain tissue and worm-algae symbiotic system, including raw and processed volumes, will be made publicly available on Zenodo upon paper acceptance. The HSAEC dataset will be available upon request.

## Code availability

All code developed for this project is available at https://github.com/Licoplion1438/Cryo-FIB-SEM

## Acknowledgments

This research is supported by NIH grant R01GM150905 (M.C.), R35GM158094 (M.X.), R01AI158569 (S.E.), R01AG082147 (I.C.) and U.S. DOE under BCIS FWP-101359 (M.C.). S.E. is a Chan Zuckerberg Biohub investigator. M.M. was supported by the Stanford Pandemic Preparedness Hub Fellowship.

## Author contributions

N.Z.: Methodology, Software, Validation, Formal analysis, Investigation, Visualization, Writing - Original Draft, Writing - Review and Editing. M.C.: Conceptualization, Methodology, Data Curation, Supervision, Funding acquisition, Writing - Review and Editing. L.J., G.W., G.S., P.L., M.M., P.L., J.S.: Resources. L.J., G.W., G.S.: Writing - Review and Editing. S.E., B.W., I.C., M.X.: Supervision, Funding acquisition.

## Competing interests

The authors declare no competing interests.

## Supplementary information

**Supplementary Fig. 1.**
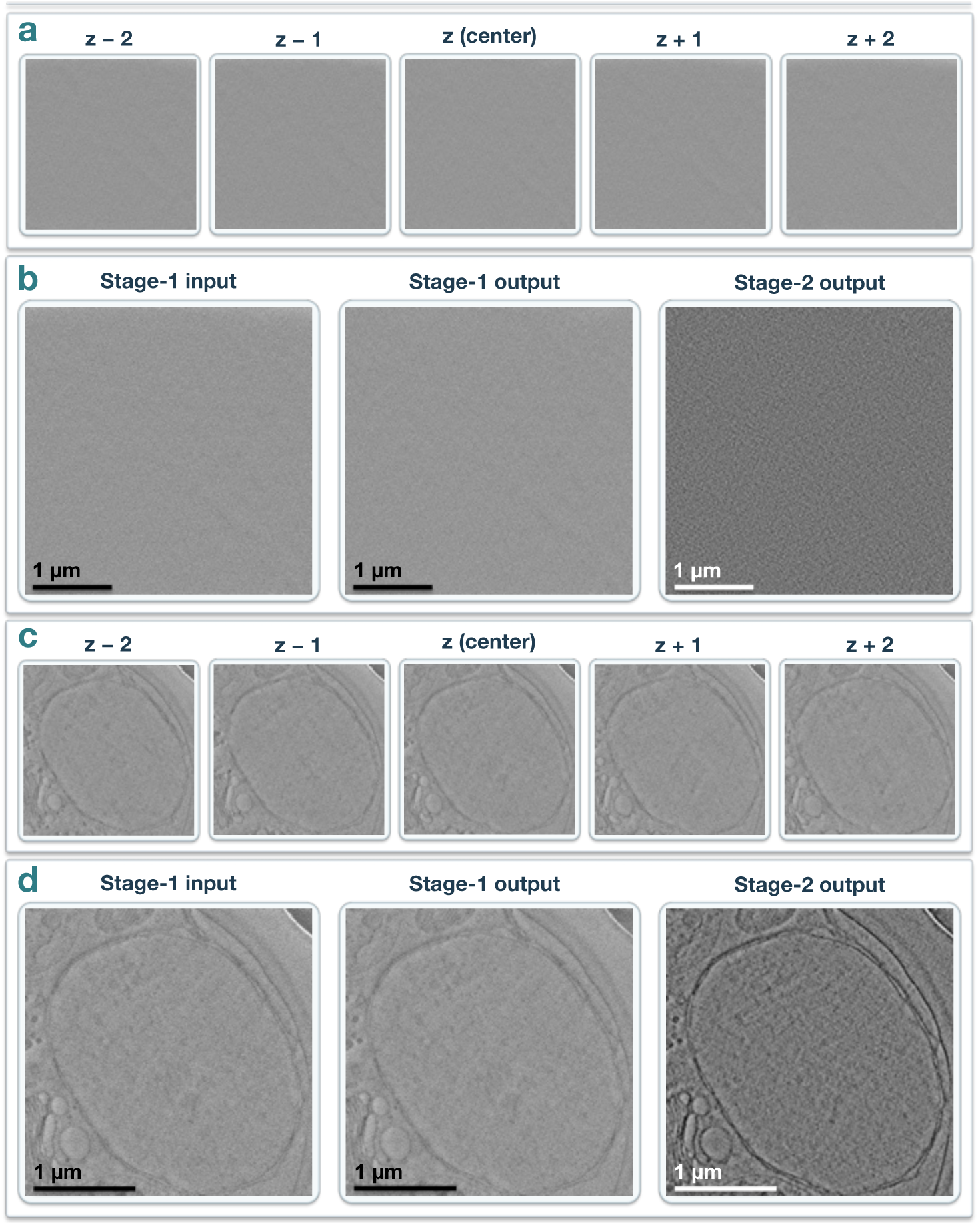
Assessment of spurious membrane generation in structurally sparse regions **a,b,** A structure-free ice region from the publicly available EMPIAR-11275 *Chlamydomonas* cryo-FIB-SEM dataset. **a,** Five consecutive Stage-1 input slices. **b,** Stage-1 input, Stage-1 output and Stage-2 output for the corresponding center slice. **c,d,** The cell nucleus region with a visible enclosing boundary but no internal membrane structure. **c,** Five consecutive Stage-1 input slices. **d,** The corresponding center-slice input and two restoration outputs. The displayed Stage-1 slices provide an overview of the available five-slice context, rather than the preprocessed and locally aligned 256 *×* 256-pixel patches actually passed to the Stage-2 network. No conspicuous new membrane-like boundaries are evident within the displayed structure-poor areas; these examples are qualitative checks, not a comprehensive assessment of spurious structure generation. Scale bars, 1 *µ*m.

**Supplementary Fig. 2.**
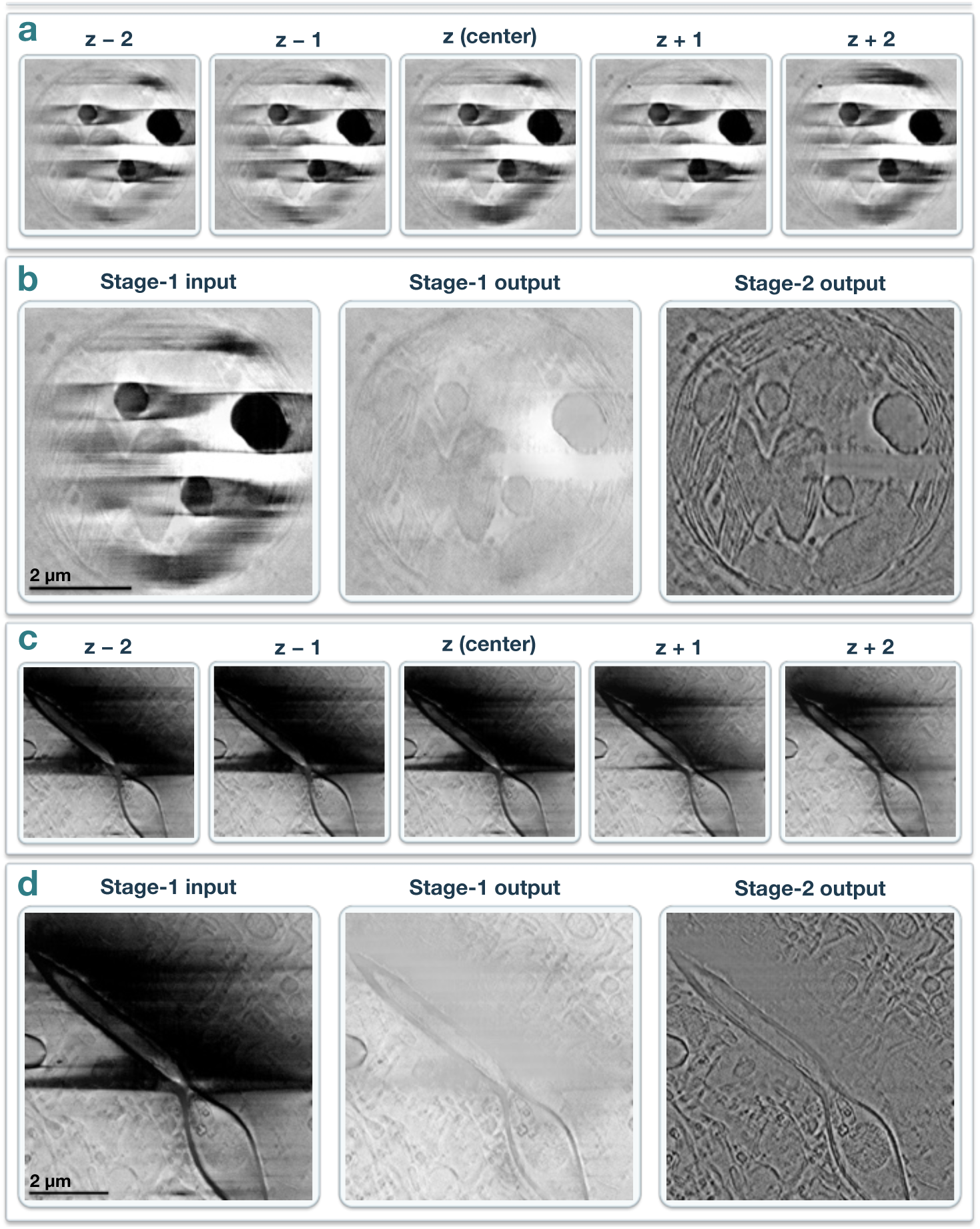
Membrane inference under varying cross-slice information loss Two-stage processing of regions with missing structural information. a,b, Worm-algae symbiotic system, illustrating persistent information loss across consecutive slices. **a,** Five consecutive Stage-1 input slices showing a region with no discernible structural information throughout the five-slice range. **b,** Stage-1 input, Stage-1 output and Stage-2 output for the center slice, showing that no apparent membrane structures emerge within the persistently obscured region. **c,d,** Mouse brain, illustrating information loss that varies across neighboring slices. **c,** Five consecutive Stage-1 input slices showing a region with no discernible structural information in the center slice, while partial structural information becomes visible in the following slice. **d,** Corresponding center-slice images showing that the two-stage processing reveals partial membrane structures within the obscured region. Scale bars, 2 *µ*m (b,d).

**Supplementary Fig. 3.**
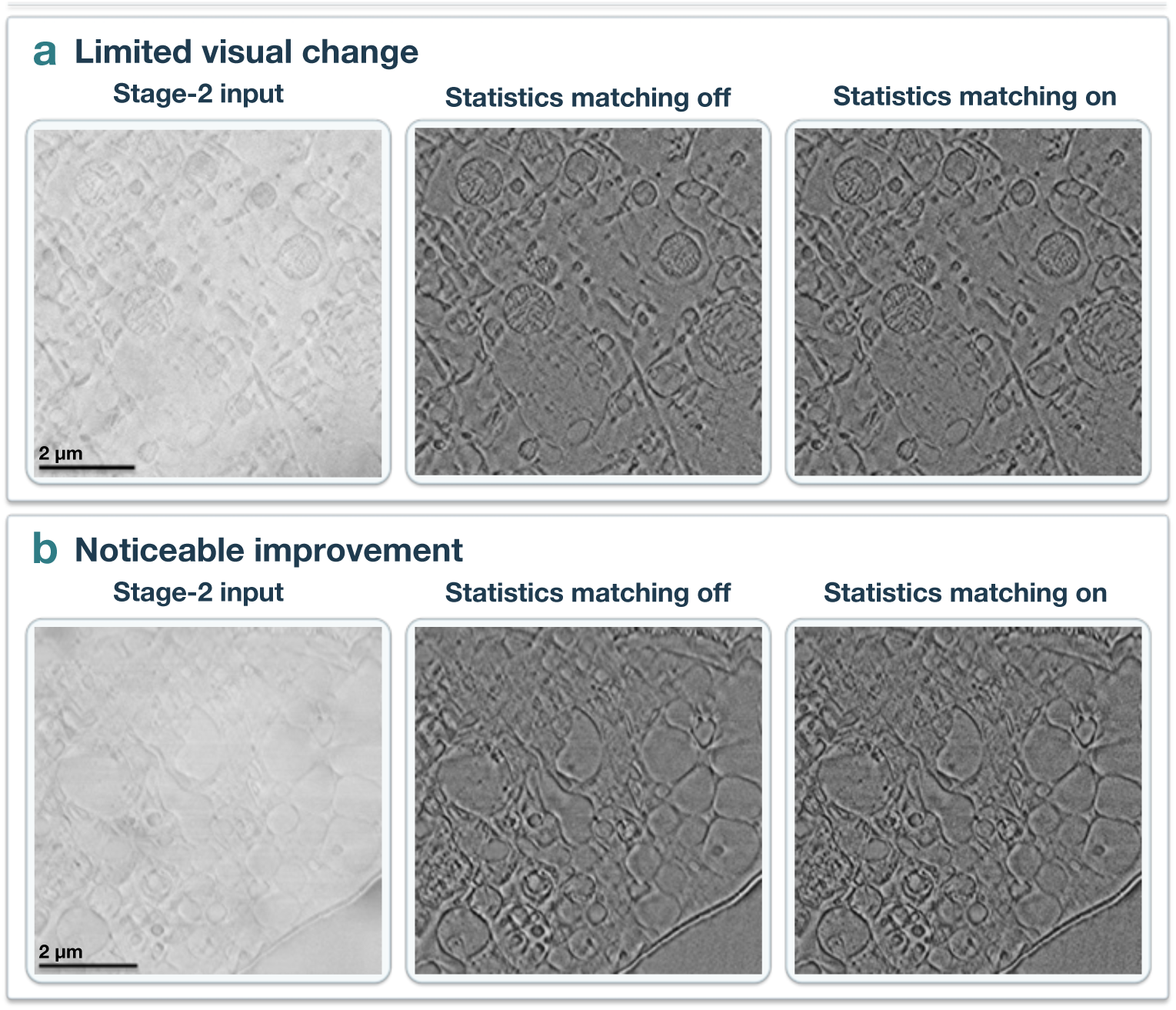
Optional Stage-2 statistics matching **Effect of optional intensity-and Fourier-statistics matching during Stage-2 inference. a,** Mouse brain, showing minimal visual differences with and without intensity matching. **b,** HSAEC, where enabling intensity matching produces slightly more pronounced membrane contrast enhancement in some regions, while the overall structural appearance remains largely unchanged. For each sample, the three columns show the Stage-2 input, Stage-2 output without intensity matching and Stage-2 output with intensity matching (left to right). Both outputs were generated using the same trained Stage-2 model. Scale bars, 2 *µ*m (left column, applicable to all images in each row).

**Supplementary Fig. 4.**
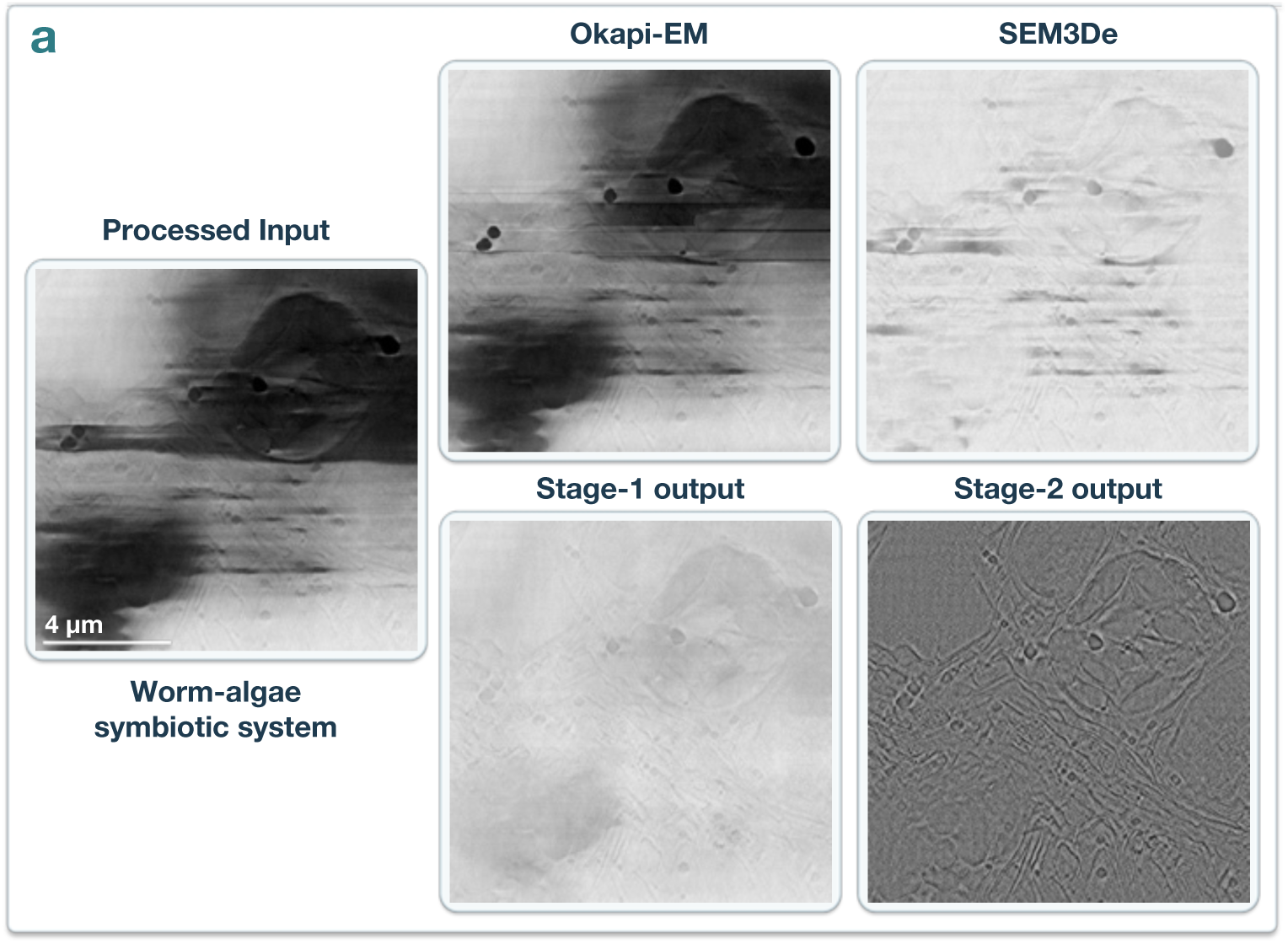
Baseline Comparison Comparison with established charge-correction methods on experimental cryo-FIB-SEM data. a, Representative region of the worm-algae symbiotic system containing overlapping charging artifacts, directional streaks and spatially varying background intensity. The large image on the left shows the processed input. The upper-right images show the same field after correction with Okapi-EM (Chafer) and SEM3De (charge-effect correction), respectively. Okapi-EM was applied with a manually annotated charging-core mask (nlinesaverage=40, minimum fitting length 50 pixels and maximum fitting edge length 700 pixels). SEM3De used rolling-ball background subtraction (radius 50 pixels, with *Light background* enabled and smoothing disabled). The lower-right images show the Stage-1 output and the subsequent Stage-2 output of our framework. Stage-1 provides the direct comparison with the artifact-correction baselines, while Stage-2 additionally enhances membrane contrast. Scale bar, 4 *µ*m (shown in the processed input and applicable to all images).

**Supplementary Fig. 5.**
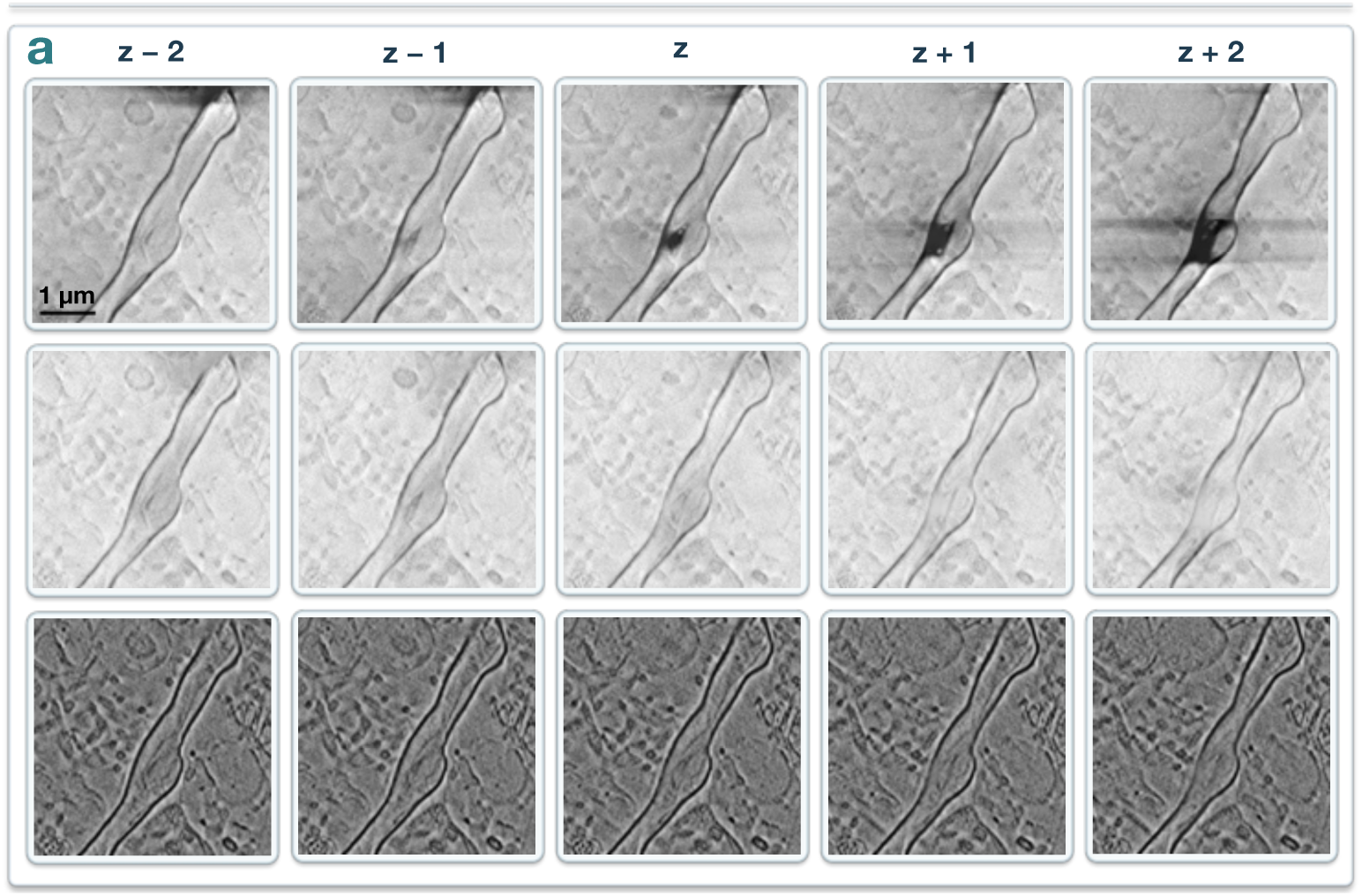
Cross-slice structural consistency Cross-slice structural consistency in mouse brain cryo-FIB-SEM data. a, Corresponding regions from five consecutive slices (*z −* 2 to *z* + 2) of a mouse brain volume. The top, middle and bottom rows show the Stage-1 input, Stage-1 output and Stage-2 output, respectively. Membrane contours exhibit consistent spatial trajectories across adjacent slices following artifact suppression and subsequent membrane contrast enhancement. Scale bar, 1 *µ*m (top-left image; applies to all images).

**Supplementary Fig. 6.**
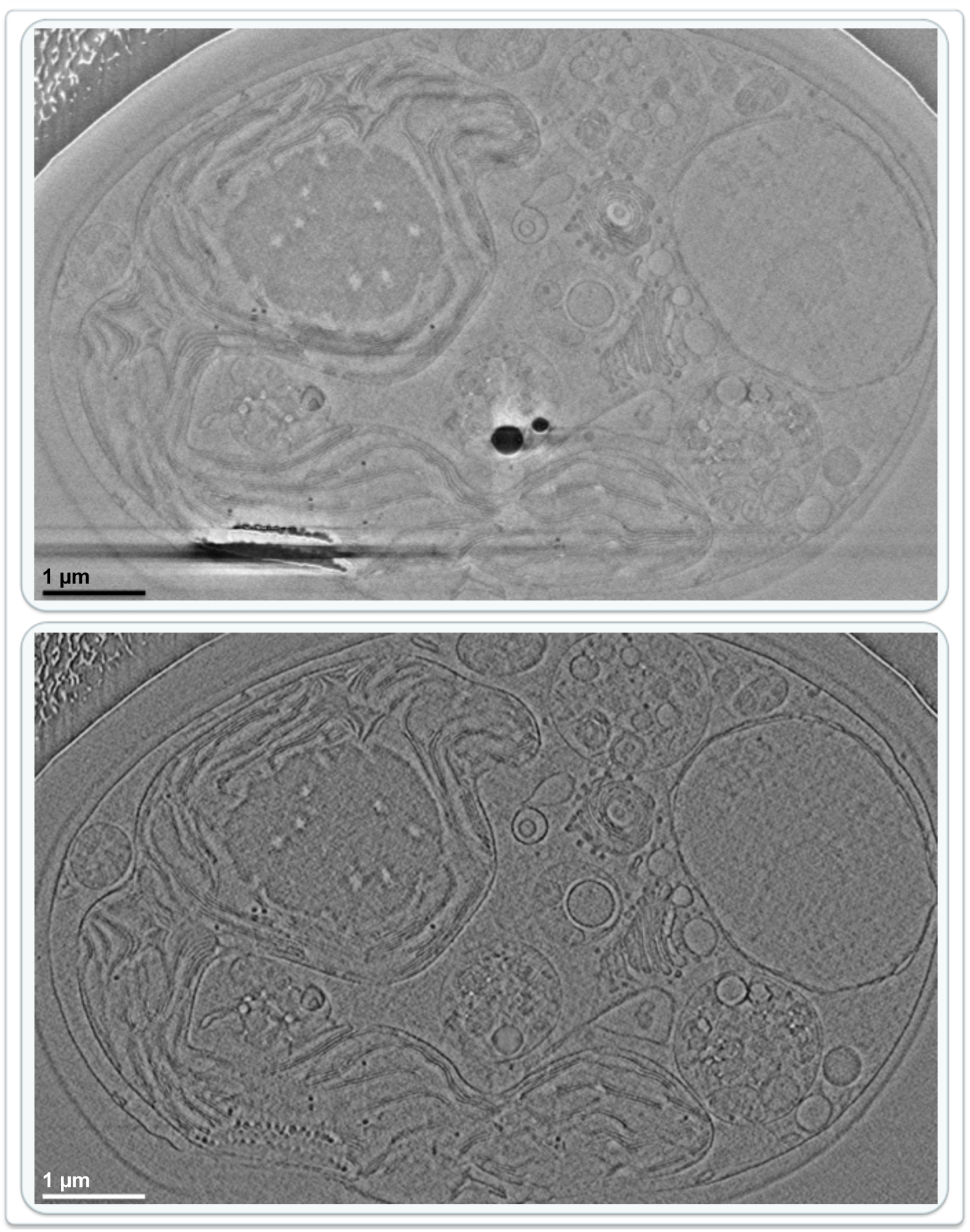
Chlamydomonas reinhardtii Larger-field view of *Chlamydomonas reinhardtii* cryo-FIB-SEM data. The images show a larger field of view from the same slice presented in Fig. 4a (EMPIAR-11275). The upper image shows the Stage-1 input and the lower image shows the corresponding Stage-2 output after two-stage processing. Scale bars, 1 *µ*m (both images).

**Supplementary Fig. 7.**
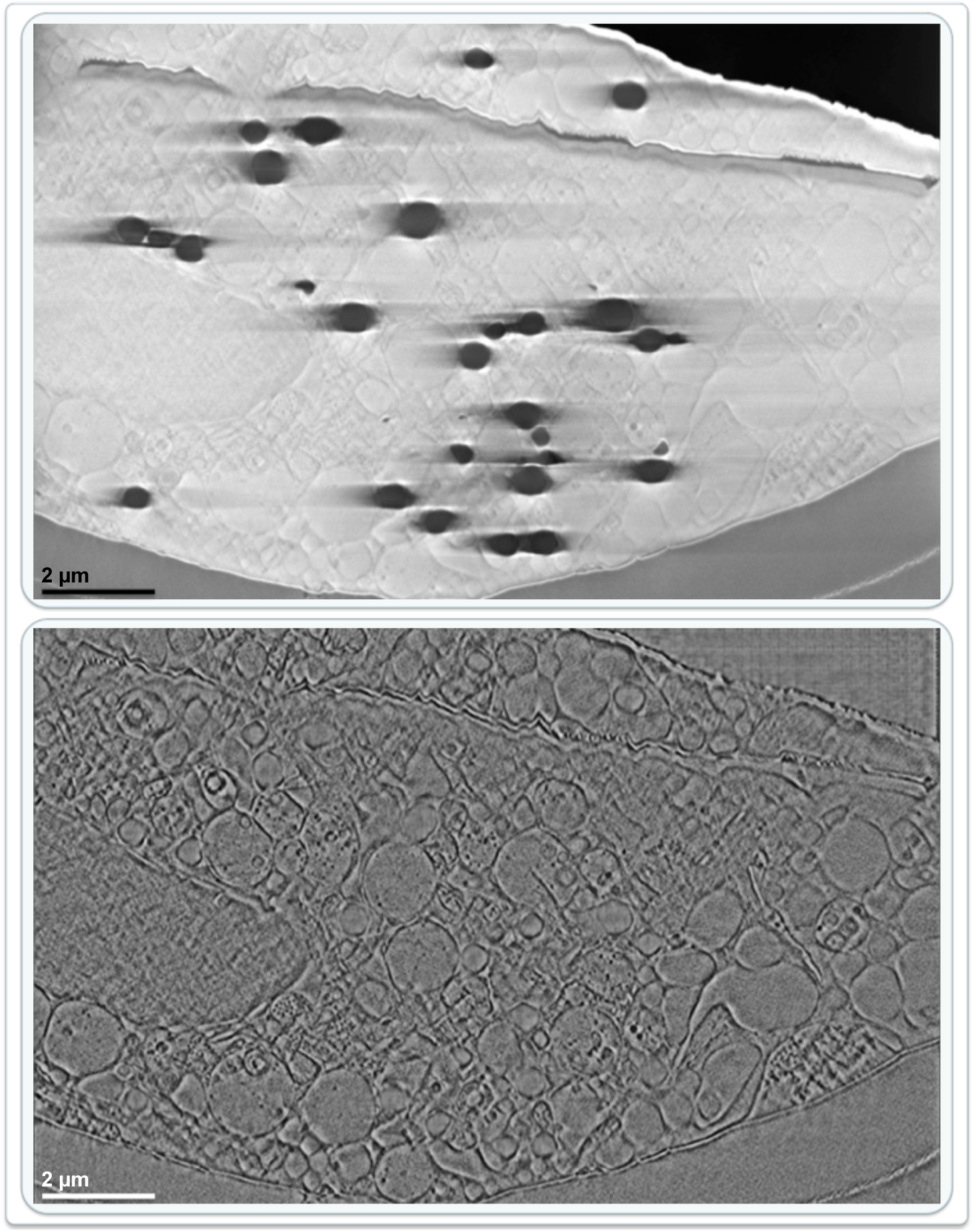
HSAEC Larger-field view of primary human small airway epithelial cells (HSAEC). The images show a larger field of view from the same slice presented in Fig. 4b. The upper image shows the Stage-1 input with charging-related intensity distortions, and the lower image shows the corresponding Stage-2 output after two-stage processing. Scale bars, 2 *µ*m (both images).

**Supplementary Fig. 8.**
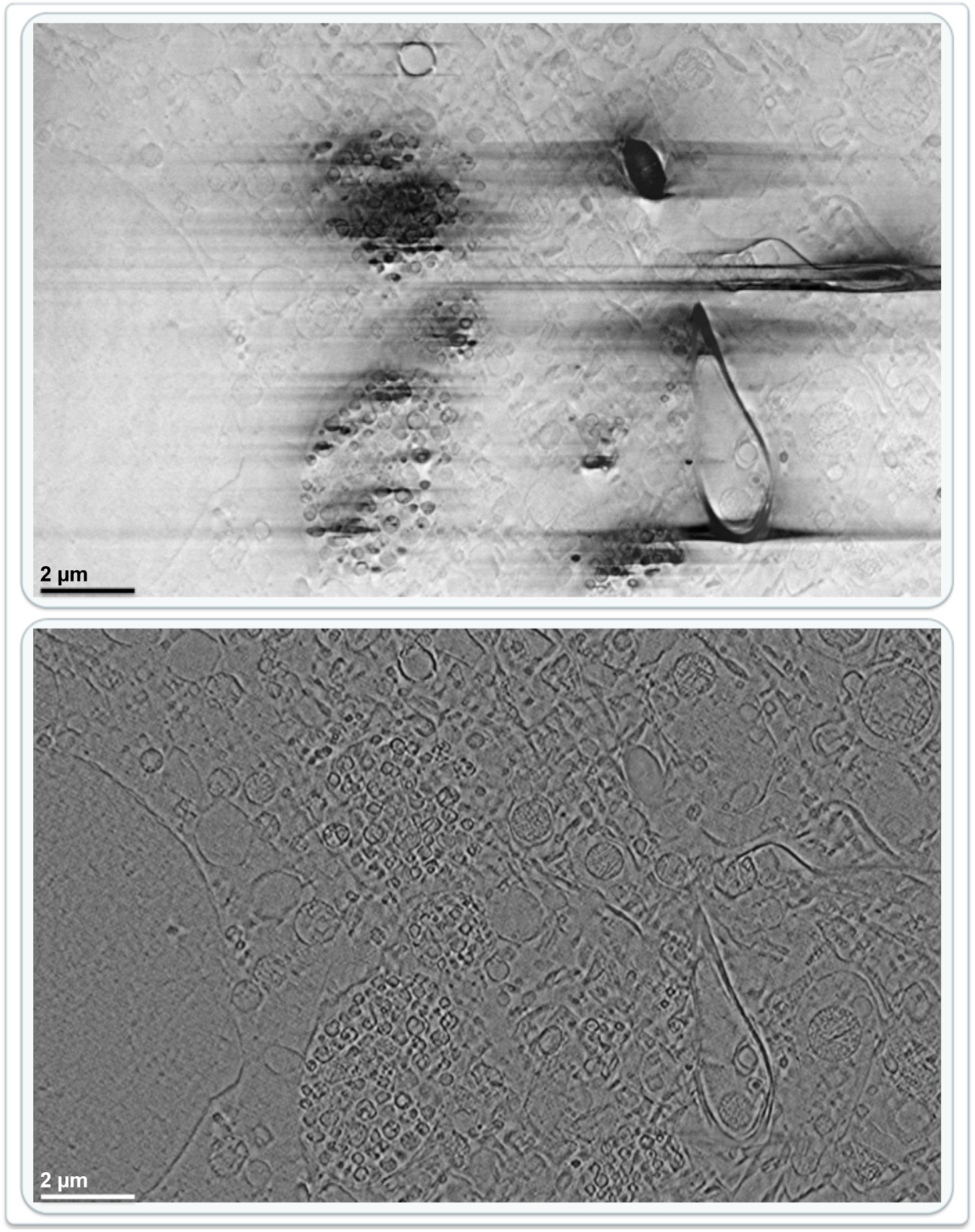
Mouse brain Larger-field view of mouse brain cryo-FIB-SEM data. The images show a larger field of view from the same slice presented in Fig. 4c. The upper image shows the Stage-1 input with localized charging and stripe artifacts, and the lower image shows the corresponding Stage-2 output after two-stage processing. Scale bars, 2 *µ*m (both images).

**Supplementary Fig. 9.**
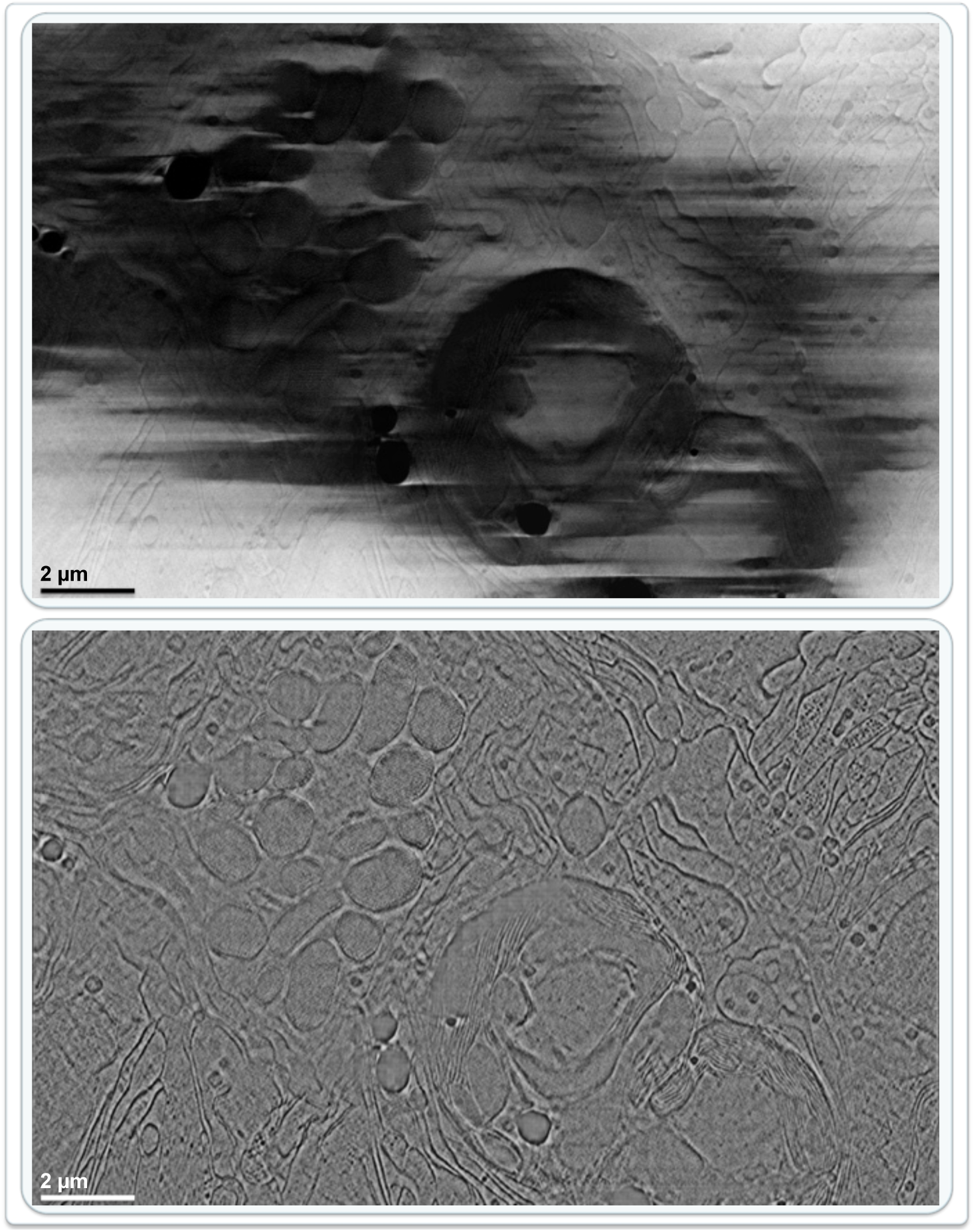
Worm-algae symbiotic system Larger-field view of the worm–algae symbiotic system. The images show a larger field of view from the same slice presented in Fig. 4d. The upper image shows the Stage-1 input with overlapping charging artifacts and spatially varying intensity, and the lower image shows the corresponding Stage-2 output after two-stage processing. Scale bars, 2 *µ*m (both images).

## Notes

### Competing Interest Statement

The authors have declared no competing interest.

## References

[1] Xu, C. S. et al. Enhanced FIB-SEM systems for large-volume 3D imaging. eLife 6, e25916 (2017).

[2] Xu, C. S. et al. An open-access volume electron microscopy atlas of whole cells and tissues. Nature 599, 147–151 (2021).

[3] Heinrich, L. et al. Whole-cell organelle segmentation in volume electron microscopy. Nature 599, 141–146 (2021).

[4] Spehner, D. et al. Cryo-FIB-SEM as a promising tool for localizing proteins in 3D. Journal of Structural Biology 211, 107528 (2020).

[5] Dumoux, M. et al. Cryo-plasma FIB/SEM volume imaging of biological specimens. eLife 12, e83623 (2023).

[6] de Beer, M. et al. Precise targeting for 3D cryo-correlative light and electron microscopy volume imaging of tissues using a FinderTOP. Communications Biology 6, 510 (2023).

[7] Velazco, A. et al. Reduction of SEM charging artefacts in native cryogenic biological samples. Nature Communications 16, 5204 (2025).

[8] Perdigão, L. M. A. et al. Okapi-EM: A napari plugin for processing and analyzing cryogenic serial focused ion beam/scanning electron microscopy images. Biological Imaging 3, e9 (2023).

[9] Serir, R. H. et al. SEM3De: image restoration for FIB-SEM. Bioinformatics Advances 3, vbad119 (2023).

[10] Lu, C. et al. Diffusion-based deep learning method for augmenting ultrastruc-tural imaging and volume electron microscopy. Nature Communications 15, 4677 (2024).

[11] González-Ruiz, V. & Fernández, J.-J. Motion-corrected Noise2Noise denoising for structural analysis of cells and tissues with (cryo-)FIB/SEM. Bioinformatics 42, btag765 (2026).

[12] Hennies, J. et al. CebraEM: A practical workflow to segment cellular organelles in volume SEM datasets using a transferable CNN-based membrane prediction (2023). URL 10.1101/2023.04.06.535829. bioRxiv:2023.04.06.535829.

[13] Chen, L., Chu, X., Zhang, X. & Sun, J. Simple baselines for image restoration, Lecture Notes in Computer Science, 17–33 (Springer, 2022).

[14] Ronneberger, O., Fischer, P. & Brox, T. U-net: Convolutional networks for biomedical image segmentation, Vol. 9351 of Lecture Notes in Computer Science, 234–241 (Springer, 2015).

[15] Fehrenbach, J., Weiss, P. & Lorenzo, C. Variational algorithms to remove stationary noise: Applications to microscopy imaging. IEEE Transactions on Image Processing 21, 4420–4430 (2012).

[16] Kelley, R., Khavnekar, S., Wietrzynski, W., Plitzko, J. & Kotecha, A. Chlamydomonas Cryo-Slice and View on Thermo Scientific Helios 5 Hydra PFIB. Electron Microscopy Public Image Archive, accession EMPIAR-11275 (2022). URL https://www.ebi.ac.uk/empiar/EMPIAR-11275/. Publicly archived cryo-FIB-SEM image dataset.

[17] Xia, D. et al. Novel App knock-in mouse model shows key features of amyloid pathology and reveals profound metabolic dysregulation of microglia. Molecular Neurodegeneration 17, 41 (2022).

[18] Nanes Sarfati, D., et al. Coordinated wound responses in a regenerative animal-algal holobiont. Nature Communications 15, 4032 (2024).

[19] Dorkenwald, S. et al. Automated synaptic connectivity inference for volume electron microscopy. Nature Methods 14, 435–442 (2017).

[20] Hennies, J. CebraNET cellular membranes in volume SEM (2023). URL 10.5281/zenodo.10047209.

[21] Schaffer, M. et al. A cryo-FIB lift-out technique enables molecular-resolution cryo-ET within native caenorhabditis elegans tissue. Nat. Methods 16, 757–762 (2019).

[22] Loshchilov, I. & Hutter, F. Decoupled weight decay regularization (2019). URL https://openreview.net/forum?id=Bkg6RiCqY7.

[23] Tomasi, C. & Manduchi, R. Bilateral filtering for gray and color images, 839–846 (1998).

[24] Srivastava, N., Hinton, G., Krizhevsky, A., Sutskever, I. & Salakhutdinov, R. Dropout: A simple way to prevent neural networks from overfitting. Journal of Machine Learning Research 15, 1929–1958 (2014). URL https://jmlr.org/papers/v15/srivastava14a.html.

[25] LeCun, Y. A., Bottou, L., Orr, G. B. & Müller, K.-R. in Efficient backprop 2 edn, Vol. 7700 of Lecture Notes in Computer Science 9–48 (Springer, 2012).

[26] Kingma, D. P. & Ba, J. Adam: A method for stochastic optimization (2015). URL https://arxiv.org/abs/1412.6980. arXiv:1412.6980.

[27] Reddy, B. S. & Chatterji, B. N. An FFT-based technique for translation, rotation, and scale-invariant image registration. IEEE Transactions on Image Processing 5, 1266–1271 (1996).

[28] Yang, Y. & Soatto, S. FDA: Fourier domain adaptation for semantic segmentation, 4085–4095 (2020).

[29] Lewis, J. P. Fast template matching, 120–123 (1995).

[30] Wang, Z., Bovik, A. C., Sheikh, H. R. & Simoncelli, E. P. Image quality assessment: from error visibility to structural similarity. IEEE Transactions on Image Processing 13, 600–612 (2004).

[31] Pettersen, E. F. et al. UCSF ChimeraX: Structure visualization for researchers, educators, and developers. Protein Science 30, 70–82 (2021).

